# FADS2-mediated fatty acid desaturation and cholesterol esterification are signatures of metabolic reprogramming during melanoma progression

**DOI:** 10.1101/2020.07.12.198903

**Authors:** Hyeon Jeong Lee, Zhicong Chen, Marianne Collard, Jiaji G Chen, Muzhou Wu, Rhoda M Alani, Ji-Xin Cheng

**Affiliations:** Photonics Center, Boston University, Boston, MA 02215, USA; College of Biomedical Engineering and Instrument Science, Zhejiang University, Hangzhou 310027, China; Key Laboratory for Biomedical Engineering of Ministry of Education, Zhejiang University, Hangzhou 310027, China; Department of Electrical and Computer Engineering, Boston University, Boston, MA 02215, USA; Department of Urology, Peking University First Hospital, Beijing 100034, China; Department of Dermatology, Boston University School of Medicine, Boston, MA 02118, USA; Department of Biomedical Engineering, Boston University, Boston, MA 02215, USA; These authors contributed equally: Hyeon Jeong Lee, Zhicong Chen

## Abstract

Identifying metabolic alterations in disease progression has been challenged by difficulties in tracking metabolites at sub-cellular level. Here, by high-resolution stimulated Raman scattering and pump-probe imaging and spectral phasor analysis of melanoma cells grouped by MITF/AXL expression pattern and of human patient tissues paired by primary and metastatic status, we identify a metabolic switch from a pigment-containing phenotype in low-grade melanoma to a lipid-rich phenotype in metastatic melanoma. The lipids found in MITF^low^/AXL^high^ melanoma cells contain high levels of cholesteryl ester (CE) and unsaturated fatty acid species. Elevated fatty acid uptake activity in MITF^low^/AXL^high^ melanoma contributes to the lipid-rich phenotype, and inhibiting fatty acid uptake suppresses cell migration. Importantly, monounsaturated sapienate is identified as an essential fatty acid that effectively promotes cancer migration. Blocking either FADS2-mediated lipid desaturation or SOAT-mediated cholesterol esterification effectively suppresses the migration capacity of melanoma *in vitro* and *in vivo*, indicating the therapeutic potential of targeting these metabolic pathways in metastatic melanoma. Collectively, our results reveal metabolic reprogramming during melanoma progression, and highlight metabolic signatures that could serve as targets for metastatic melanoma treatment and diagnosis.

## Main

Melanoma is the most aggressive form of skin cancer. When melanoma is caught in its early stages and surgically removed, the prognosis is favorable; once melanoma has metastasized, it becomes difficult to treat ^1,2^. Although increasing Breslow depth, or melanoma tumor thickness, is associated with worse survival ^3^, the majority of deaths attributed to melanoma are due to thin tumors (<1 mm) ^4^. With the incidence of melanoma steadily rising at a rate of 3% per year ^5^, it is more important than ever to detect melanoma at an early stage and identify the tumors that are the most at-risk for metastasizing for early intervention. Nevertheless, without molecular signatures, precision diagnosis and treatment of metastatic melanoma remains difficult.

One promising target is melanin, the major pigment in melanoma. It was found that more eumelanin than pheomelanin is present in malignant melanoma ^6,7^. A recent study used pump-probe microscopy as an elegant label-free approach to distinguish the two types of melanin in melanoma tissues ^8^, heralding potential prognostic significance ^9^. Despite this advance, loss of microphthalmia-associated transcription factor (MITF), the gene regulating pigmentation, is reported in melanomas with an invasive phenotype ^10,11^, indicating an unmet need to identify new molecular markers for detecting aggressive and invasive melanoma.

Melanoma exhibits a dynamic and adaptive phenotypic switching capability ^12^. Among the signaling pathways mediating such process, MITF status is considered a central regulator of the phenotypic switch ^13^, namely from a highly proliferative, less invasive to a less proliferative, highly invasive state. Specifically, loss of MITF is an indication of de-differentiation of melanoma ^14,15^, which is supported by enhanced migration/invasion gene expression profiles ^16^. Despite of these advances in genetic characterization of melanoma phenotypes, metabolic markers for advanced melanoma have just begun to be explored. A recent report suggested that elevated lipid uptake in metastatic melanoma ^17,18^, indicating that cancer progression is partly sustained by high lipogenic or lipid uptake activity; however, due to difficulties in compositional analysis, the exact metabolites and precise mechanisms that regulate melanoma progression remains unknown.

Here, we report findings of metabolic reprogramming in metastatic melanoma by multimodal chemical imaging tools recently developed. Specifically, by stimulated Raman scattering (SRS) imaging of biomolecules ^19,20^ and pump-probe imaging of pigments ^21^ at a sub-cellular level, we revealed a previously unrecognized metabolic switch from pigmentation in low-grade melanoma to lipid droplet (LD) accumulation in metastatic melanoma. Raman spectroscopic analysis further identified a significant amount of unsaturated fatty acids and cholesteryl ester (CE) in these LDs. Blocking fatty acid uptake and depleting LDs dramatically suppresses cell migration. In particular, one unsaturated fatty acid, sapienate, synthesized through fatty acid desaturase 2 (FADS2), is found to play an essential role in promoting melanoma migration through regulation of membrane fluidity. Mass spectrometry further confirms a higher level of sapienate in lipid-rich, metastatic melanoma compared to lipid-poor, non-invasive melanoma. Consequently, inhibition of FADS2 suppresses melanoma migration *in vitro* and metastasis *in vivo*. In parallel, inhibiting cholesterol esterification is found to significantly reduce LD accumulation and suppress cell migration by inactivation of the Wnt/β-catenin pathway. Together, these results present a unique metabolic reprogramming profile and provide diagnostic signatures and therapeutic targets for metastatic melanoma.

## Results

### Melanoma progression into metastatic state is accompanied by a metabolic switch from pigmentation to lipid accumulation

To investigate the metabolic reprogramming during melanoma progression, human melanoma cell lines bearing high MITF or low MITF expression were used. Consistent with previous reports ^11,22^, the melanoma cells with high MITF expression levels exhibit low AXL Receptor Tyrosine Kinase (AXL) expression (**Supplementary Fig. 1**). On the basis of this expression profile, the human melanoma cell lines used in this study (**Supplementary Table 1**) were clustered into MITF^high^/AXL^low^ or MITF^low^/AXL^high^ groups. To map the various metabolites in human melanoma cells, we deployed stimulated Raman scattering microscopy ^19,20^ to map chemical species at sub-cellular resolution. The laser beating frequency was initially tuned to be resonant with C-H stretching vibration at 2899 cm^−1^. We detected a stimulated Raman loss signal arising from C-H rich biomolecules, showing a clear cell morphology with a slightly weaker signal from the nucleus. We observed a number of droplets in melanocytes, MITF^high^/AXL^low^, and MITF^low^/AXL^high^ melanoma cells (**Fig. 1a**). These droplet structures in an SRS image are often considered to be lipid droplets (LDs) ^19^. To verify the chemical specificity, we tuned the laser beating frequency to 2265 cm^−1^, off-resonant with C-H stretching vibration. No signal was observed from the off-resonance image of MITF^low^/AXL^high^ melanoma cells, which suggests that these droplets are rich in C-H bonds and likely are LDs (**Fig. 1a**). This assignment was confirmed by immunofluorescence staining of a LD-associated protein, adipophilin (**Supplementary Fig. 2**). Intriguingly, droplets with strong signals were still observed in melanocytes and MITF^high^/AXL^low^ melanoma, indicating that signals from these droplets are not specific to C-H stretching vibration (**Fig. 1a**). We attribute the strong signal at off-Raman resonance to either transient absorption or photothermal effects, which is often detected with a pump-probe imaging scheme ^23,24^.

**Figure 1.**
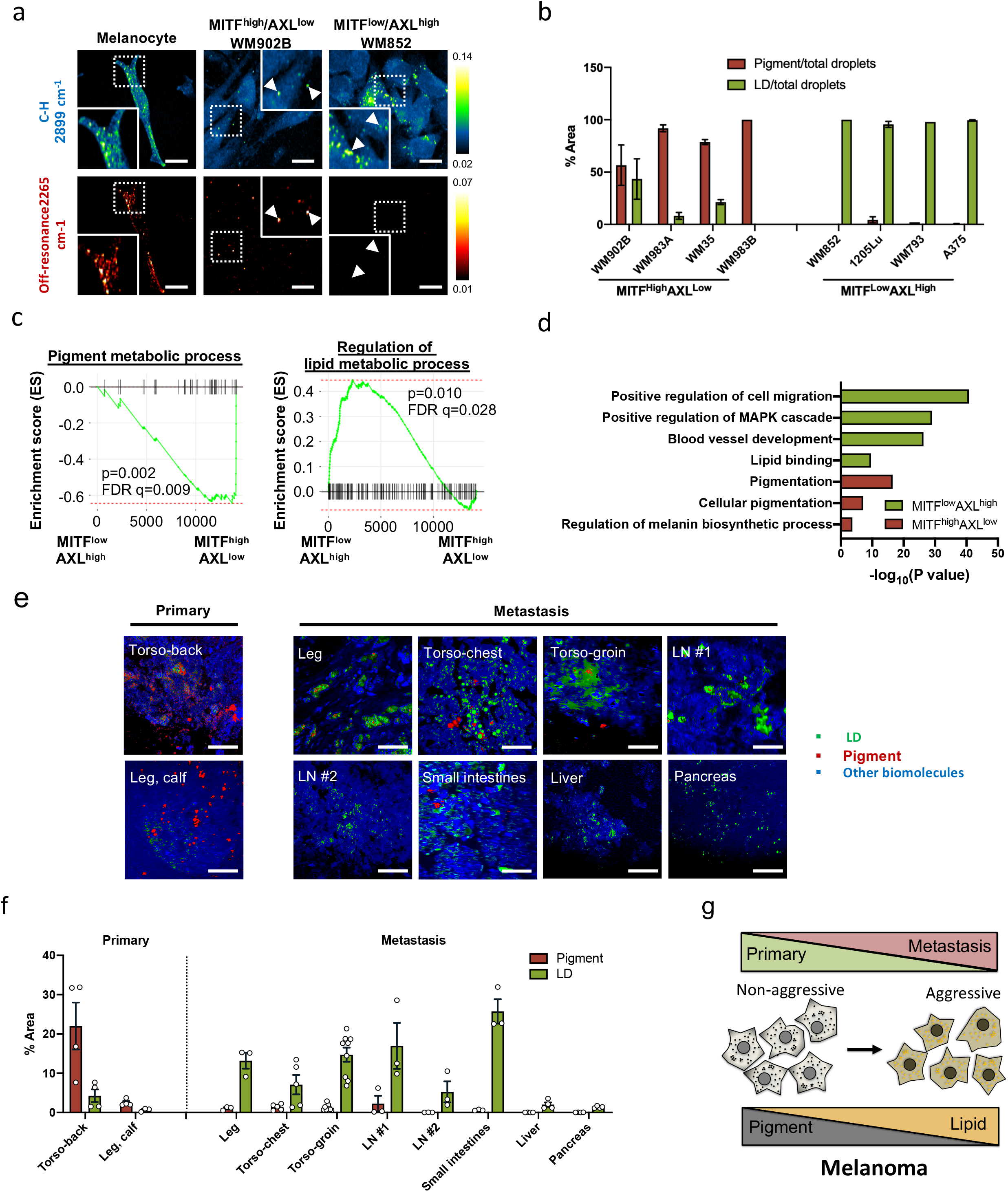
Metastatic melanoma exhibits LD-rich phenotype. (a) Representative SRS images of human melanoma cells in the C-H region (2899 cm^−1^) and in the off-resonance region (2265 cm^−1^). Arrows indicate droplets. Scale bars, 20 μm. (b) Quantification of droplets identified as pigment and LD from phasor analysis. (c) GSEA of human melanoma samples from TCGA-SKMC based on MITF and AXL expression profile. (d) Functional enrichment network analysis for MITF^low^/AXL^high^ melanoma and MITF^high^/AXL^low^ melanoma. (e) Multimodal imaging and phasor output of patient primary and metastatic melanoma tissue samples. LN, lymph node; SI, small intestines. Scale bars, 50 μm. (f) Percent area of pigment and LD presented in the tissue. (g) Metabolic switching observed during melanoma progression from primary to metastasis. Data represent mean ± standard error of the mean (SEM).

To verify this hypothesis, we performed a time-resolved measurement. The signals from these droplets show a fast decay within 1.0 picosecond and then remained at the same level (**Supplementary Fig. 3a**). The fast decay and the remaining signal are attributed to transient absorption and photothermal processes, respectively ^25,26^. In comparison, the pump-probe decay curve of eumelanin powder extracted from *Sepia officinalis* showed a similar profile (**Supplementary Fig. 3b**). Unlike the pigments in melanoma, the pigments from melanocytes exhibit a negative decay curve (**Supplementary Fig. 3a**), indicating a different type of pigments presented in melanocytes. It was reported that when imaged under a pump-probe microscope, pheomelanin exhibits a negative signal whereas eumelanin exhibits a positive signal ^8^. Thus, it is likely that the pigments in melanoma are mostly eumelanin and the pigments in melanocytes are mostly pheomelanin. Consistent with this observation, it was reported that eumelanin is presented in higher content in melanoma compared to nonmalignant nevi ^6,8^. Collectively, these results support that both melanocytes and MITF^high^/AXL^low^ melanoma cells have high pigmentation activity due to high MITF expression.

To establish a method for quantifying pigmentation and LD accumulation in melanoma, we designed a time-domain multimodal SRS/pump-probe imaging and phasor analysis approach (**Supplementary Fig. 4**). In SRS imaging, 802 nm serves as a pump beam and 1045 nm serves as a Stokes beam, while in pump-probe imaging, 1045 nm serves as a pump beam and 802 nm serves as a probe beam (**Supplementary Fig. 4a**). SRS and pump-probe signals show different profiles in the time-domain: SRS displays a sharp gaussian-shape peak, while pump-probe displays a slow decay curve (**Supplementary Fig. 4b, c**). Therefore, by performing time-resolved measurement followed by phasor analysis ^27^, these two signals can be well-separated (**Supplementary Fig. 4d**). We found that the subtle SRS spectral difference between LDs versus rest of the cell can be further separated through phasor analysis (**Supplementary Fig. 4d**). Through the multimodal imaging and phasor analysis approach, we simultaneously imaged LD accumulation and pigmentation in human melanoma cell lines and quantified the amounts of LDs and pigments in the samples (**Fig. 1b, Supplementary Fig. 4e, f**). There is a high percentage of pigment droplets in MITF^high^/AXL^low^ melanoma. Nevertheless, almost all droplets identified in MITF^low^/AXL^high^ melanoma are LDs (**Fig. 1b**). Together, these data indicate a reduction of pigmentation and increase of lipid accumulation during the progression of melanoma from low grade to high grade.

Consistent with the metabolic profile, the gene set enrichment analysis (GSEA) of human melanoma samples from The Cancer Genome Atlas (TCGA) showed that the pigment metabolic process gene set was enriched in MITF^high^/AXL^low^ group, while regulation of lipid metabolic process gene set was enriched in MITF^low^/AXL^high^ group (**Fig. 1c**, **Supplementary Table 2**). Differentially expressed genes (DEGs) were generated base on the median expression level of MITF and AXL (**Supplementary Fig. 5a**, **Supplementary Table 3**). Functional enrichment network analysis for MITF^low^/AXL^high^ melanoma identified several gene sets related to aspects of cancer metastasis such as cell migration and blood vessel development, and lipid binding process that were enriched from the up-regulated genes, while gene sets related to pigmentation were enriched from the down-regulated genes (**Fig. 1d**, **Supplementary Fig. 5b**, **Supplementary Table 3**). These results indicate a distinct metabolic profile of MITF^high^/AXL^low^ versus MITF^low^/AXL^high^ melanoma.

To validate the clinical relevance of such metabolic switch in metastatic melanoma, we performed multimodal imaging of human primary and metastatic melanoma tissue samples (**Fig. 1e**). The tumor lesions were confirmed by pathological assessment of the neighboring slices stained with hematoxylin and eosin (**Supplementary Fig. 6**). By time-resolved nonlinear optical imaging and phasor analysis, signals from pigment, LDs, and other biomolecules were separated and quantified (**Fig. 1f**). Primary melanoma tissues contain higher pigment content than LD content. In contrast, metastatic melanoma, including metastatic lesions on the extreme leg, torso-chest, and torso-groin, and metastatic tumor in other organs such as lymph nodes, small intestines, liver, and pancreas, contain higher LD content than pigment content (**Fig. 1f**). On the basis of these results, we hypothesize that there is a metabolic switch from pigmentation to lipid accumulation during the progression of human melanoma from the primary tumor to the metastatic tumor (**Fig. 1g**).

### The LDs in MITF^low^/AXL^high^ melanoma are composed of CE and unsaturated lipids

To unveil the composition of LDs accumulated in MITF^low^/AXL^high^ melanoma, we performed confocal Raman spectroscopic measurement of single LDs (**Fig. 2a**). The Raman spectra of LDs in all MITF^low^/AXL^high^ melanoma cells showed similar spectral profiles, including bands for lipids in the fingerprint region (1200 – 1800 cm^−1^) and prominent CH_2_ stretching band (2850 cm^−1^). More detailed spectral analysis revealed that these LDs show peaks from the C=C stretching vibration at 1654 cm^−1^ and from the vibration of =C-H bonds at 3002 cm^−1^, both suggesting the presence of unsaturated lipids (**Fig. 2a**). To quantify the unsaturation degree, we generated a calibration curve using fatty acids containing various numbers of C=C bonds (**Supplementary Fig. 7a**). Raman spectra of palmitate (no C=C bond), oleate (one C=C bond) and linoleate (two C=C bonds) were obtained and analyzed. The peak intensity of C=C stretching vibration at 1654 cm^−1^ is found to be linearly correlated with the number of C=C bonds present in the fatty acids after normalization by the CH_2_ bending band at 1445 cm^−1^ (**Supplementary Fig. 7a,b**). On the basis of this calibration, we estimated the unsaturation degree to be 1.2 in WM852, 1.3 in 1205Lu, 1.4 in WM793, and 1.8 in A375 cells (**Fig. 2b**). To further validate the presence of unsaturated lipids in LDs, we treated LD-rich melanoma cells with desaturase inhibitors, CAY10566 and SC26196, for inhibiting SCD and FADS2, respectively. Cells treated with either SCD or FADS2 inhibitor showed reduced peak intensities at both 1654 cm^−1^ and 3002 cm^−1^ (**Fig. 2c**). The quantification of desaturation degree further showed the reduction of desaturation degree to below or close to 1 after treatment with desaturase inhibitors (**Fig. 2d**), supporting the reduction of unsaturated lipids. Together, these results indicate that the LDs accumulated in MITF^low^/AXL^high^ melanoma contain a significant amount of unsaturated fatty acids.

**Figure 2.**
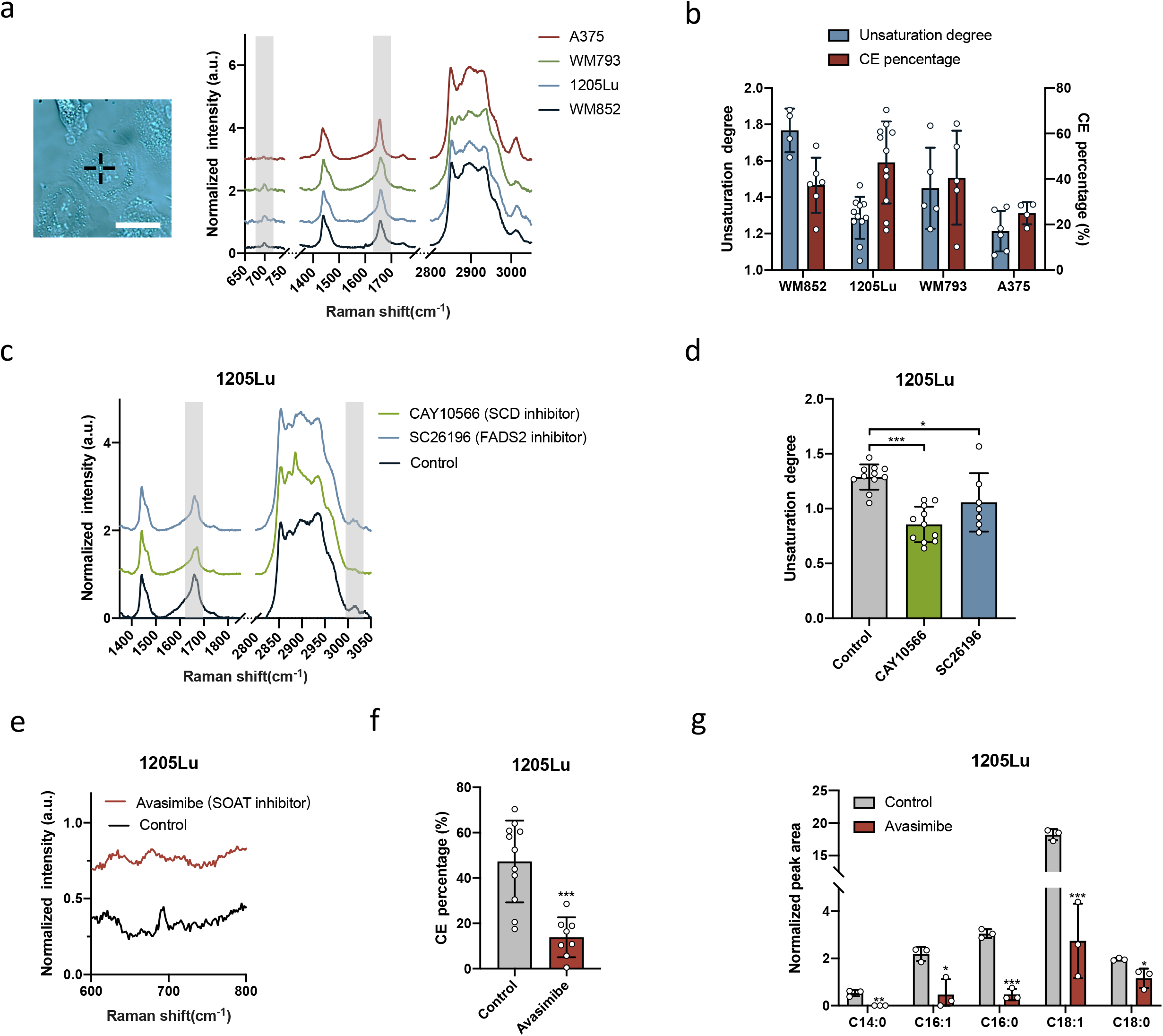
LDs in aggressive melanoma are composed of unsaturated fatty acids and CE. (a) Representative Raman spectra of single LDs in MITF^low^/AXL^high^ melanoma. Spectral intensity was normalized by the peak at 1445 cm^−1^. Grey areas highlight the Raman peaks used to quantify unsaturation degree and CE percentage. Insert shows the brightfield image of single LD identified in a cell. Scale bar: 10 μm. (b) Quantification of unsaturation degree and CE percentage based on the calibration curves. (c) Representative Raman spectra of LDs in 1205Lu treated with DMSO as control, CAY10566 (50 μM, 2 days) and SC26196 (50 μM, 2 days). Grey areas highlight the Raman peaks indicative of unsaturation lipids. (d) Quantification of unsaturation degree calculated from the calibration curve. (e) Representative Raman spectra of LDs in 1205Lu treated with DMSO as control and avasimibe (10 μM, 2 days). (f) Quantification of CE percentage in LDs calculated from the calibration curve. (g) Quantification of LC/MS measurement of lipids extracted from control and avasimibe-treated 1205Lu cells (10 μM, 2 days). Data represent mean ± SD. *: p < 0.05, **: p < 0.01, ***: p < 0.001.

Another important component that we identified from the Raman spectral profile is cholesteryl ester (CE). This feature was indicated by the cholesterol band at 702 cm^−1^ and the C=O ester stretching band at 1740 cm^−1^ (**Fig. 2a**). To quantify the CE percentage, we generated calibration curves for CE percentage using CE and triacylglycerol emulsions (**Supplementary Fig. 7c,d**). The Raman spectra of lipid emulsions with various ratios of CE and triacylglycerol showed that the peak intensity of cholesterol band at 702 cm^−1^ was linearly correlated to the molar percentage of CE in the lipid emulsions after normalization by the CH_2_ bending band at 1445 cm^−1^ (**Supplementary Fig. 7c,d**). On the basis of the calibration, it was estimated that the CE contents of the LDs are 37.3%, 47.3%, 40.6%, and 25.0% in WM852, 1205Lu, WM793, and A375, respectively (**Fig. 2b**). Inside cells, excess cholesterol is known to be esterified by the sterol O-acyltransferase (SOAT) enzyme and stored in LDs ^28^. To validate that CE is stored in LDs, we treated cells with a potent SOAT inhibitor, avasimibe. After avasimibe treatment, the peak intensity at 702 cm^−1^ reduced significantly (**Fig. 2e**). The quantification of CE percentage in LDs showed that LDs are composed of less than 20% CE when cholesterol esterification is inhibited by avasimibe (**Fig. 2f**). Furthermore, the LC/MS measurement of extracted lipids from MITF^low^/AXL^high^ melanoma cells confirmed the CE-rich lipid profile and significant reduction of multiple CE species after avasimibe treatment (**Fig. 2g**, **Supplementary Fig. 7e**). Moreover, the mass spectrometry data identified cholesteryl oleate (CE 18:1) to be the dominant species. Together, these results indicate that LDs accumulated in MITF^low^/AXL^high^ melanoma are comprised of unsaturated fatty acids and CE.

### Fatty acid uptake is the major pathway for LD accumulation in MITF^low^/AXL^high^ melanoma

There are two major routes for lipid accumulation in mammalian cells: one is *de novo* lipogenesis, and the other is fatty acid uptake through transporter proteins (**Fig. 3a**). In *de novo* lipogenesis, glucose most often serves as the precursor, from which the acetyl-CoA is generated and used for lipid synthesis. Fatty acid uptake is facilitated by various fatty acid transporters, such as FABP4, CD36, and FATPs ^29^. In both cases, excess fatty acids are stored in LDs. To understand how MITF^low^/AXL^high^ melanoma cells obtain the lipids stored in the LD, we performed SRS imaging of deuterated metabolites in melanoma cells to evaluate their glucose-derived lipogenesis and fatty acid uptake activities. As SRS imaging of Glucose-D_7_ has been used to track the *de novo* lipogenesis activity of cancer cells ^30^, we cultured MITF^high^/AXL^low^ and MITF^low^/AXL^high^ melanoma cells with a glucose-D_7_-containing medium. SRS imaging in the C-D vibration at 2127 cm^−1^ was performed to visualize glucose-D_7_-derived metabolites (**Fig. 3b**). Overall, a weak signal was observed from the cytoplasm in both MITF^high^/AXL^low^ and MITF^low^/AXL^high^ melanoma cells (**Fig. 3b**). No obvious droplet structures were observed in both groups. The quantification of intracellular C-D signal showed no significant difference between the two groups (**Fig. 3c**), indicating no significant difference in glucose-derived metabolites, including *de novo* lipogenesis.

**Figure 3.**
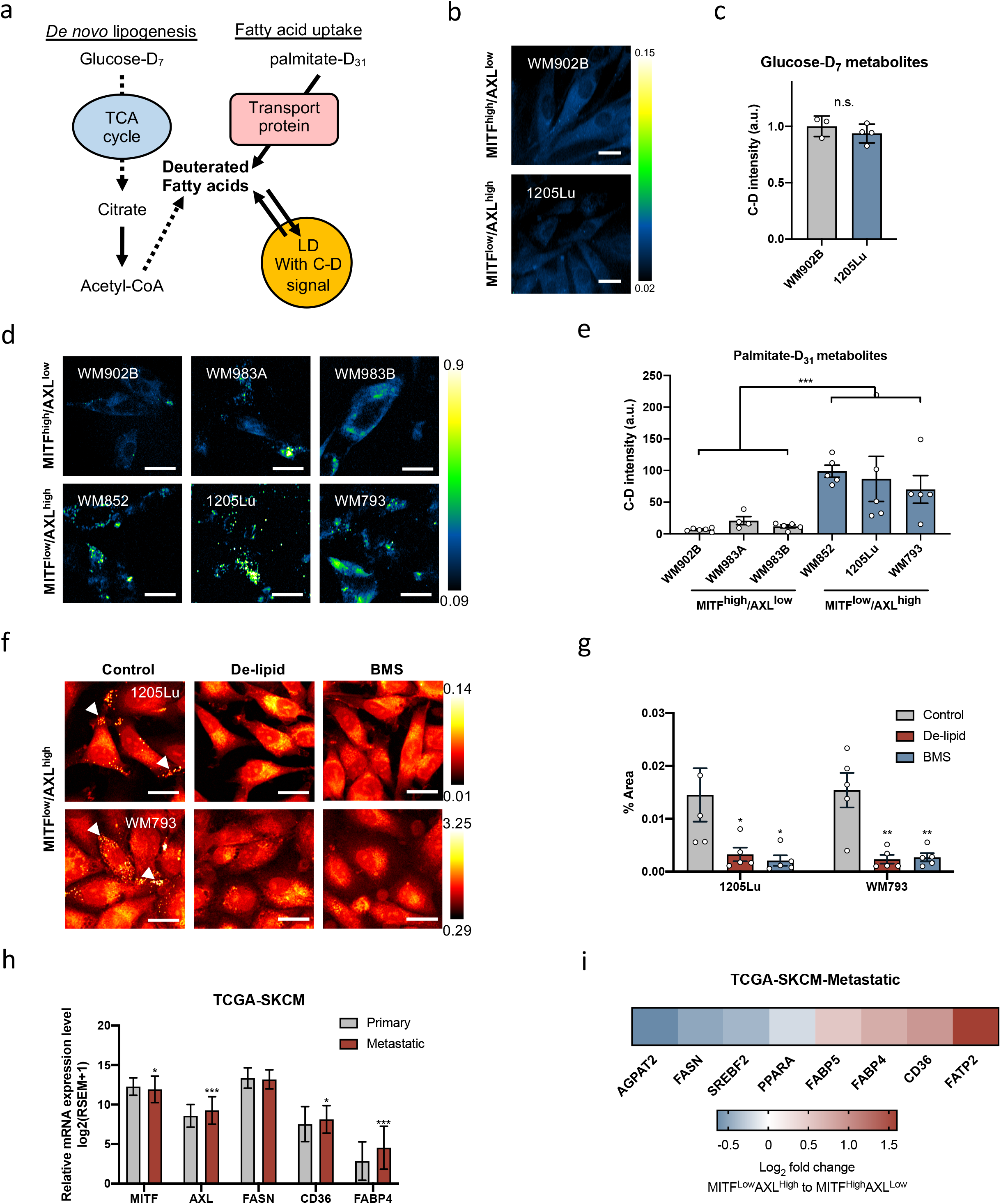
Fatty acid uptake is the major source for LD accumulation in MITF^low^/AXL^high^ melanoma. (a) A schematic of lipid sources contributing to LD synthesis and deuterated metabolite tracking method. (b) Representative SRS images in the C-D region (2127 cm^−1^) of WM902B and 1205Lu, cultured with glucose-D_7_ containing media for 2 days. (c) Quantification of SRS intensity at 2127 cm^−1^ from C-D positive LDs. (d) Representative SRS images in the C-D region (2127 cm^−1^) of WM902B, WM983A, WM983B, WM852, 1205Lu and WM793, cultured with palmitate-D_31_ containing media (100 μM, 6 hours). (e) Quantification of SRS intensity at 2127 cm^−1^ for C-D positive LDs. Data represent mean ± SEM. (f) Representative SRS images in the C-H region (2899 cm^−1^) of 1205Lu and WM793, cultured in normal media as control, in delipidized serum media (2 days) and treated with BMS (50 μM, 1 day). (g) Percentage area of LDs in 1205Lu and WM793 as shown in (f). (h) Relative mRNA expression levels of MITF, AXL, FASN, CD36 and FABP4 in primary and metastatic melanoma from TCGA-SKCM database. (i) Differences in expression levels of genes related to lipid metabolism between MITF^low^/AXL^high^ and MITF^high^/AXL^low^ metastatic melanoma from TCGA-SKCM database. Scale bars, 10 μm. Data represent mean ± SD unless indicated otherwise. *: p < 0.05, **: p < 0.01, ***: p < 0.001.

To evaluate fatty acid uptake activity, palmitate-D_31_ was supplemented into the culture medium. SRS imaging at 2127 cm^−1^ clearly showed LDs incorporated with deuterated fatty acids, especially in MITF^low^/AXL^high^ melanoma cells (**Fig. 3d**). To quantify LDs with the C-D signal, we performed particle analysis and measured the C-D intensity per cell area in each group. MITF^low^/AXL^high^ melanoma cells showed significantly higher C-D signal than MITF^high^/AXL^low^ melanoma cells (**Fig. 3e**), indicating high palmitate uptake activity in these cells. Similarly, when we supplemented oleate-D_34_ into the culture medium, MITF^low^/AXL^high^ cells showed higher oleate uptake activity than MITF^high^/AXL^low^ cells (**Supplementary Fig. 8**). To further validate that fatty acid uptake is the major source of LD accumulation in these cells, we suppressed fatty acid uptake by removing extracellular fatty acids using a de-lipidized serum. SRS imaging at C-H stretching vibration at 2899 cm^−1^ showed a significant reduction in the number of LDs when fatty acid availability became limited (**Fig. 3f,g**). Similarly, when we suppressed fatty acid uptake by treating the cells with an inhibitor (BMS309403) targeting fatty acid-binding protein 4 (FABP4), a fatty acid transporter, the number of LDs reduced significantly (**Fig. 3f,g**). Collectively, MITF^low^/AXL^high^ melanoma has higher fatty acid uptake activity, which serves as the major source of LD accumulation in these cells.

To validate these findings in clinical samples, we analyzed the expression of genes related to lipogenesis and fatty acid uptake in human melanoma samples from TCGA divided into primary and metastatic melanoma (**Fig. 3h**). MITF expression is reduced whereas AXL expression level increased in the metastatic melanoma group, validating that MITF^low^/AXL^high^ melanoma cells represent metastatic melanoma more closely. Importantly, the rate-limiting enzyme in *de novo* lipogenesis, fatty acid synthase (FASN), showed no difference between primary and metastatic melanoma (**Fig. 3h**), which is consistent with the SRS imaging results for evaluating lipogenesis activity shown in **Fig. 3b** and **3c**. However, fatty acid transporters such as CD36 and FABP4 were up-regulated in metastatic melanoma compared to primary melanoma (**Fig. 3h**), suggesting increased fatty acid uptake activity. The metastatic melanoma samples were further divided into MITF^high^/AXL^low^ and MITF^low^/AXL^high^ groups and used to compare the expression of lipid metabolism-related genes (**Fig. 3i**). The MITF^low^/AXL^high^ group showed up-regulation of multiple fatty acid transporters and down-regulation of genes involved in lipogenesis compared to the MITF^low^/AXL^high^ group. Together, these results support that while lipid *de novo* synthesis activity is at the same level, fatty acid uptake activity is elevated in human metastatic melanoma.

### Fatty acid sapienate significantly promotes cell migration

To understand the function of LD accumulation in melanoma, we separated melanoma cell lines based on their LD accumulation level (**Supplementary Table 1**). When we compared the migration capacity of cells, LD-rich melanoma cells showed significantly higher migration capacity than LD-poor melanoma cells (**Fig. 4a,b**). Since LD accumulation mainly comes from fatty acid uptake (**Fig. 3**), we tested the migration capacity of LD-rich cells after inhibition of fatty acid uptake to deplete LDs. When the LD accumulation is suppressed by depleting extracellular lipids with de-lipidized serum (**Fig. 3f,g**), the number of migrated cells reduced significantly (**Fig. 4c, Supplementary Fig. 9a**). Moreover, when the LD-rich cells were treated with various fatty acid transporters, including BMS309403 (FABP inhibitor), sulfosuccinimidyl oleate (CD36 inhibitor), and lipofermata (FATP inhibitor), migration capacity was significantly reduced (**Fig. 4c, Supplementary Fig. 9a**). These results indicate that LD accumulation, driven by fatty acid uptake, is a signature of enhanced melanoma migration potential.

**Figure 4.**
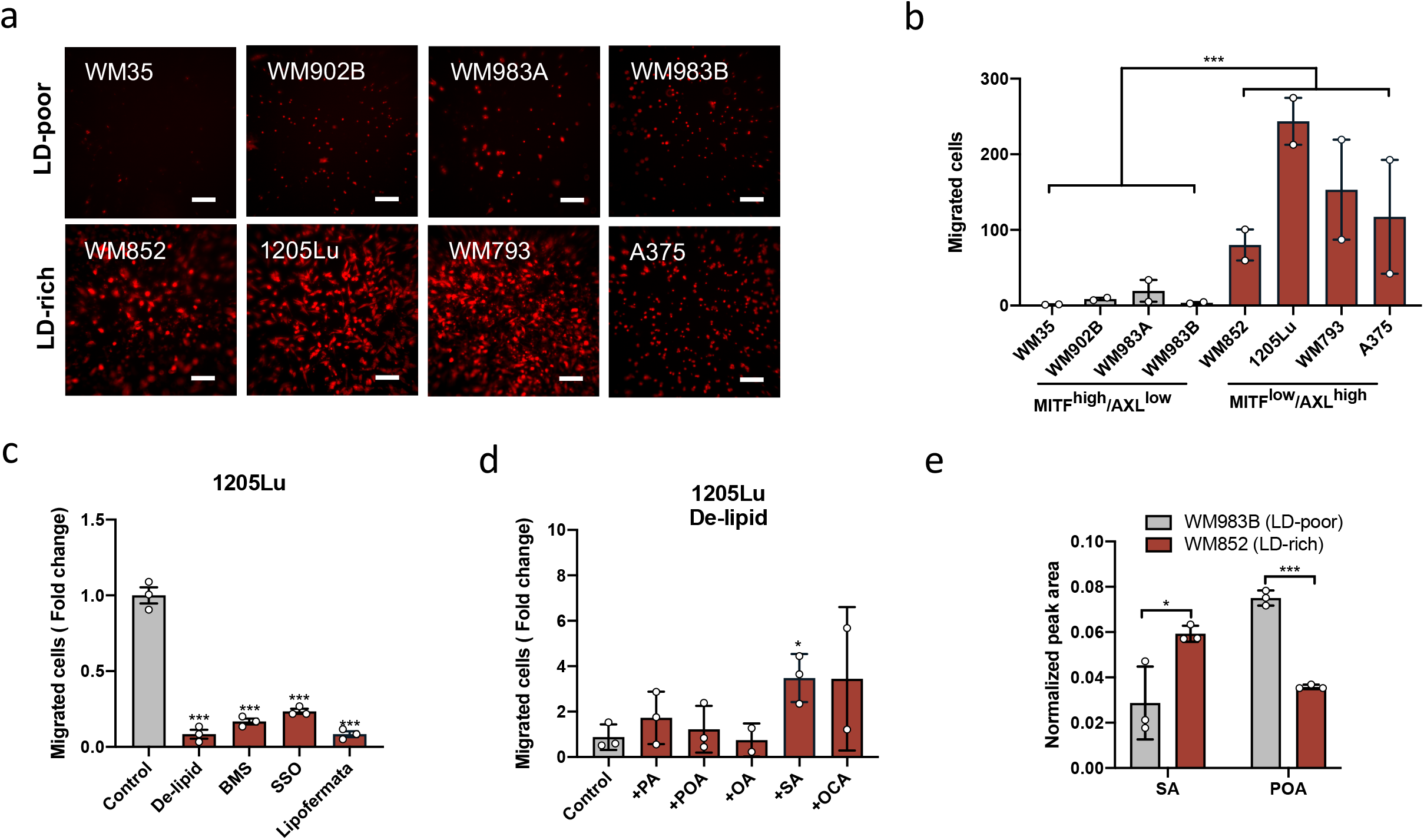
Fatty acid sapienate significantly promotes cell migration. (a) Images and (b) quantification of migrated MITF^high^/AXL^low^ and MITF^low^/AXL^high^ melanoma cells. (c) Quantification of migrated 1205Lu pre-cultured with de-lipidized medium or pre-treated with BMS309403 (BMS, 50 μM, 1 day), sulfosuccinimidyl oleate (SSO, 50 μM, 1 day) and lipofermata (10 μM, 1 day). Control group was used for normalization. (d) Quantification of migrated 1205Lu pre-cultured with de-lipidized serum media (1 day) and supplemented with ethanol as control and fatty acids (20 μM, 12 hours) as indicated. Control group was used for normalization. (e) Quantification of normalized peak areas of sapienate and palmitoleate presented in WM852 and WM983B. Scale bars, 50 μm. Data represent mean ± SEM. *: p < 0.05, ***: p < 0.001.

Next, we asked whether certain fatty acids play a more important role in promoting cell migration than others. To test this possibility, we evaluated the migration rescue capability of fatty acids by supplementing them into the de-lipidized serum (**Fig. 4d, Supplementary Fig. 9b**). Palmitate, palmitoleate, and oleate did not rescue cell migration in de-lipidized serum (**Fig. 4d, Supplementary Fig. 9b**). Intriguingly, supplementation of sapienate and its elongated form, octadecenoate, rescued cell migration significantly (**Fig. 4d, Supplementary Fig. 9b**), indicating an essential role of these fatty acids in cell migration. To validate the presence of sapienate in LD-rich cells, we measured and compared sapienate levels between LD-rich and LD-poor melanoma. Both sapienate (*cis*-6-hexadecenoate) and palmitoleate (*cis*-9-hexadecenoate) are monounsaturated fatty acids that enhance the unsaturation degree in cells, but the positions of the double bond are different. To specifically measure sapienate, we applied GC/MS to separate and detect sapienate and palmitoleate. Sapienate shows a peak at an earlier retention time than palmitoleate (**Supplementary Fig. 10a**), and this protocol allows us to separate these two fatty acids to obtain the amount of sapienate in the sample. By fatty acid methyl ester (FAME) analysis, we extracted free fatty acids from LD-rich and LD-poor melanoma cells. On the basis of the GC/MS analysis, the LD-rich melanoma is found to contain much higher amount of sapienate than the LD-poor melanoma (**Fig. 4e, Supplementary Fig. 10b**). Interestingly, the palmitoleate content shows the opposite trend, with a higher amount in LD-poor melanoma than in LD-rich melanoma (**Fig. 4e**). Together, these results suggest that sapienate is an essential fatty acid in LD-rich melanoma and has an important functional role of enhancing cell migration.

### Inhibition of FADS2 suppresses human melanoma migration *in vitro* and metastasis *in vivo*

The opposite trends of sapienate and palmitoleate levels in LD-poor and LD-rich melanoma (**Fig. 4e**) triggered us to investigate the role of sapienate metabolism in melanoma aggressiveness. Sapienate is synthesized from palmitate by FADS2, while palmitoleate is synthesized from palmitate by SCD in the endothelial curriculum (**Fig. 5a**). A recent study indicates that these two desaturases can be complementary to each other to support cancer progression ^31^. Interestingly, our TCGA database analysis indicates that while SCD is expressed at similar levels, FADS2 expression is significantly up-regulated in metastatic melanoma compared to primary melanoma (**Supplementary Fig. 11a**). To further evaluate the importance of FADS2-mediated fatty acid desaturation in melanoma metastasis, we tested the effects of FADS2 inhibition on cell migration (**Fig. 5b,c**). FADS2 was inhibited by an inhibitor, SC26196, or by shRNA (**Supplementary Fig. 11b**). After inhibition of FADS2 by SC26196, cell migration is suppressed by 1.5-fold compared to the control group (**Fig. 5b**). As an additional evidence, FADS2 knockdown by shRNA resulted in reduced cell migration by up to 2-fold, depending on the degree of knockdown efficiency (**Fig. 5c** and **Supplementary Fig. 11b**). Importantly, sapienate supplementation rescued migration capacity of the cells with either FADS2 inhibition methods (**Fig. 5b,c**), indicating that sapienate has a functional role in promoting melanoma cell migration. Furthermore, FADS2 knockdown suppressed cell invasion, another important property of cancer metastasis, by 2-fold (**Supplementary Fig. 11c,d**). To evaluate the role of FADS2 in metastasis, we used a mouse tail-vein model. Human melanoma cells stably expressing control shRNA or FADS2 shRNA were intravenously injected, and the amount of lung metastases was compared (**Fig. 5d**). Mice injected with melanoma cells containing FADS2 knockdown developed less lung metastases compared to the mice injected with control melanoma cells (**Fig. 5d**). FADS2 knockdown resulted in a 68% reduction in the fractional area of lung metastases compared to the control group (**Fig. 5e**). Together, these results indicate that FADS2 is a potential target for suppressing melanoma metastasis.

**Figure 5.**
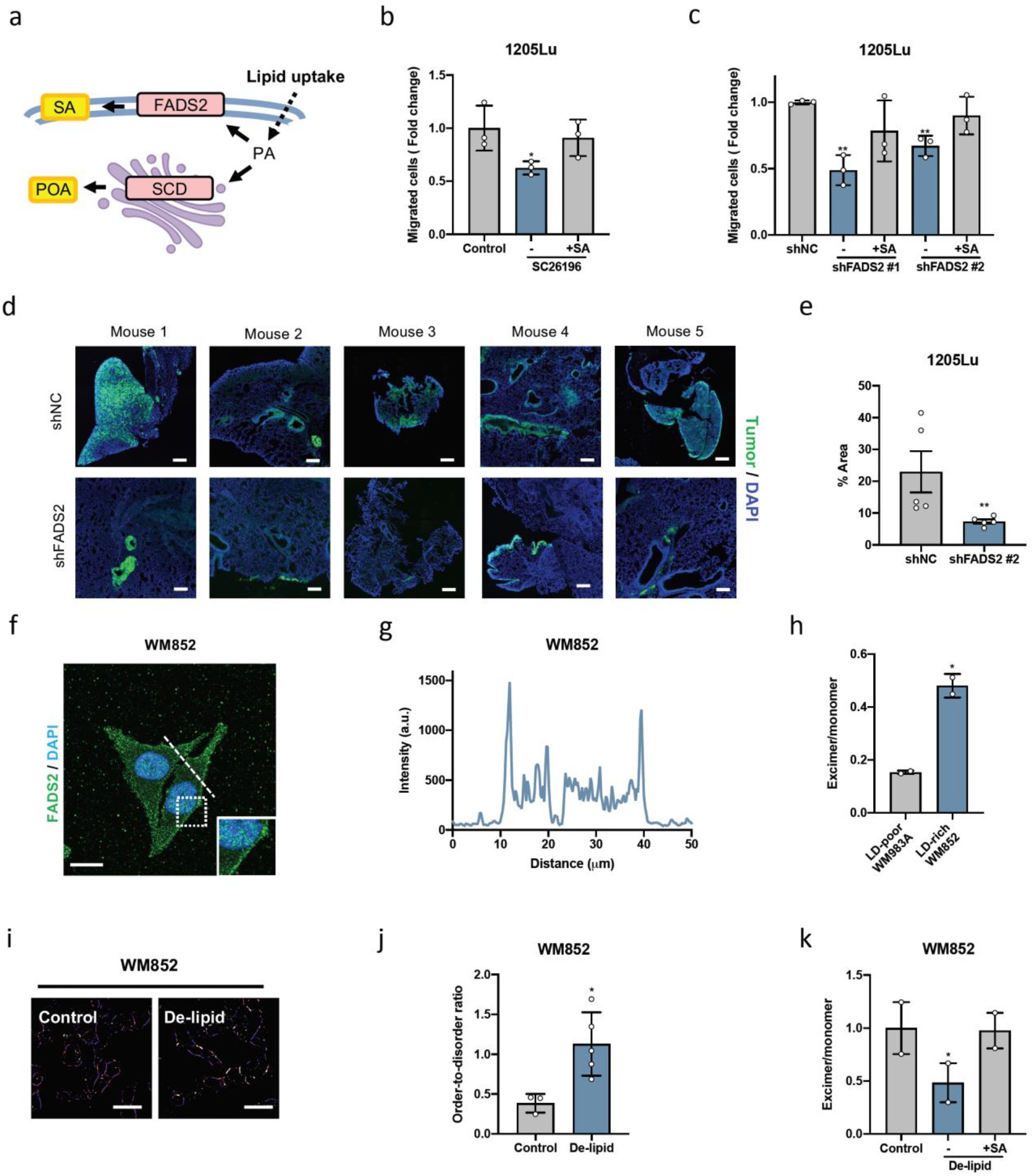
Inhibition of FADS2 suppresses melanoma migration and metastasis through modulation of membrane fluidity. (a) A schematic of two major fatty acid desaturases and their products. (b) Quantification of migrated 1205Lu treated with DMSO as a control, with SC26196 (50 μM, 2 days), and SC26196 with supplement of sapienate (20 μM, 1 day during SC26196 treatment). Control group was used for normalization. (c) Quantification of migrated 1205Lu stabling expressing control shRNA (1205Lu-shNC) or FADS2 shRNAs (1205Lu-shFADS2 #1 and #2). 1205Lu expressing FADS2 shRNAs were cultured in sapienate supplemented medium (20 μM, 1 day) for testing the rescue capability. (d) Fluorescence images of lung tissues from mice 46 days after injected with 1205Lu-shNC or 1205Lu-shFADS2 #1 expressing GFP into tail-vein. Scale bars: 200 μm. (e) Quantification of percentage area that tumors occupy in lung sections. Data represent mean ± SEM. N = 5 mice per group. (f) An image of immunofluorescence staining of FADS2 in WM852. (g) Intensity profile across the line indicated in (f). (h) The ratio between excimer and monomer of membrane fluidity probe (pyrenedecanoic acid, PDA) in WM983A and WM852. Data represent mean ± SEM. (i) Ratiometric images of membrane fluidity probe (di-4-ANESPPDHQ) in WM852 cultured in normal medium as control and de-lipidized serum medium. (j) Quantification of order-to-disorder ratio obtained from (i). (k) The ratio between excimer and monomer of PDA in WM852 cultured with normal medium as control, with de-lipidized serum medium and de-lipidized serum medium with sapienate supplement (50 μM, 1 day). Scale bars: 50 μm unless indicated otherwise. Data represent mean ± SD unlese indicated otherwise. *: p < 0.05, **: p < 0.01.

### Melanoma does not depend on fatty acid oxidation for energy production, proliferation, or migration

One of the major functions of fatty acids is providing energy through β-oxidation ^32^. To investigate whether elevated fatty acid uptake in LD-rich melanoma cells support fatty acid oxidation, we performed extracellular flux analyses using a Seahorse XFe96 analyzer on melanoma cells treated with or without 4 μM etomoxir to inhibit mitochondrial fatty acid oxidation (**Supplementary Fig. 12a,b**). There were no differences in total ATP generation between etomoxir-treated cells and control cells (**Supplementary Fig. 12a**). In more detail, we analyzed the energy metabolic status of the cells based on mitochondrial and glycolytic ATP generation (**Supplementary Fig. 12b**). There were no differences in mitochondrial or glycolytic ATP generation between etomoxir-treated cells and control cells; thus, no metabolic shift observed.

Consistent with this finding, the on-target 5 μM etomoxir dose had no effect on LD-rich melanoma cell proliferation while the off-target 200 μM etomoxir dose slightly reduced cell proliferation (**Supplementary Fig. 12c,d**). Although high doses of etomoxir are used ubiquitously, new research shows that higher doses (>10μM) have off-target effects leading to oxidative phosphorylation suppression through Complex I inhibition, which is erroneously interpreted as a reduction in fatty acid oxidation ^33,34^. Because of this, caution must be taken when interpreting data using high concentrations of etomoxir to inhibit fatty acid oxidation. Furthermore, LD-rich melanoma cells treated with the on-target 5 μM dose showed no significant change in the cell migration capacity compared to control cells (**Supplementary Fig. 12e,f**). Collectively, these results indicate that LDs in melanoma are not used for fatty acid oxidation normal, non-starved conditions.

### Sapienate promotes melanoma migration through modulation of membrane fluidity

In search of the molecular mechanism of enhanced melanoma metastasis through increased level of sapienate, we investigated the intracellular distribution of FADS2 protein by immunofluorescence (**Fig. 5f**). It is intriguing that a significant amount of FADS2 was localized on the plasma membrane (**Fig. 5f,g**). From this unique localization of FADS2 protein and the unsaturation property of its metabolic product, we hypothesize that the downstream metabolites of FADS2 modulate membrane fluidity. Increased membrane fluidity facilitates cell migration and cancer metastasis ^35,36^. Consistent with these results, LD-rich melanoma cells show higher membrane fluidity than LD-poor melanoma cells (**Fig. 5h**). Furthermore, when the fatty acid uptake is suppressed by culturing in the de-lipidized serum condition, the membrane fluidity reduced significantly by 2-fold (**Fig. 5i,j**). Importantly, sapienate supplemented into the de-lipidized serum was sufficient to rescue this effect, restoring the membrane fluidity to the level similar to the control group (**Fig. 5k**). These results suggest that sapienate promotes cell migration and cancer metastasis through increasing the membrane fluidity.

### Blocking cholesterol esterification suppresses melanoma migration through Wnt/β-catenin pathway

In addition to the finding of sapienate in melanoma, we also observed a significant amount of CE accumulation in LDs (**Fig. 2**). CE is synthesized from cholesterol and fatty acids by SOAT protein (**Fig. 6a**). To investigate the roles of cholesterol esterification in melanoma aggressiveness, we suppressed cholesterol esterification by a small molecule inhibitor avasimibe or by knockdown via shRNA (**Supplementary Fig. 13a**). When cholesterol esterification is inhibited, the number of LDs reduce significantly (**Fig. 6b,c**), suggesting a reduction in fatty acid uptake and storage. At the same time, avasimibe treatment suppressed melanoma migration significantly (**Fig. 6d, Supplementary Fig. 13b**). SOAT1 knockdown also significantly suppressed cell migration (**Fig. 6e, Supplementary Fig. 13c**). These results indicate that cholesterol esterification is another potential target for metastatic melanoma.

**Figure 6.**
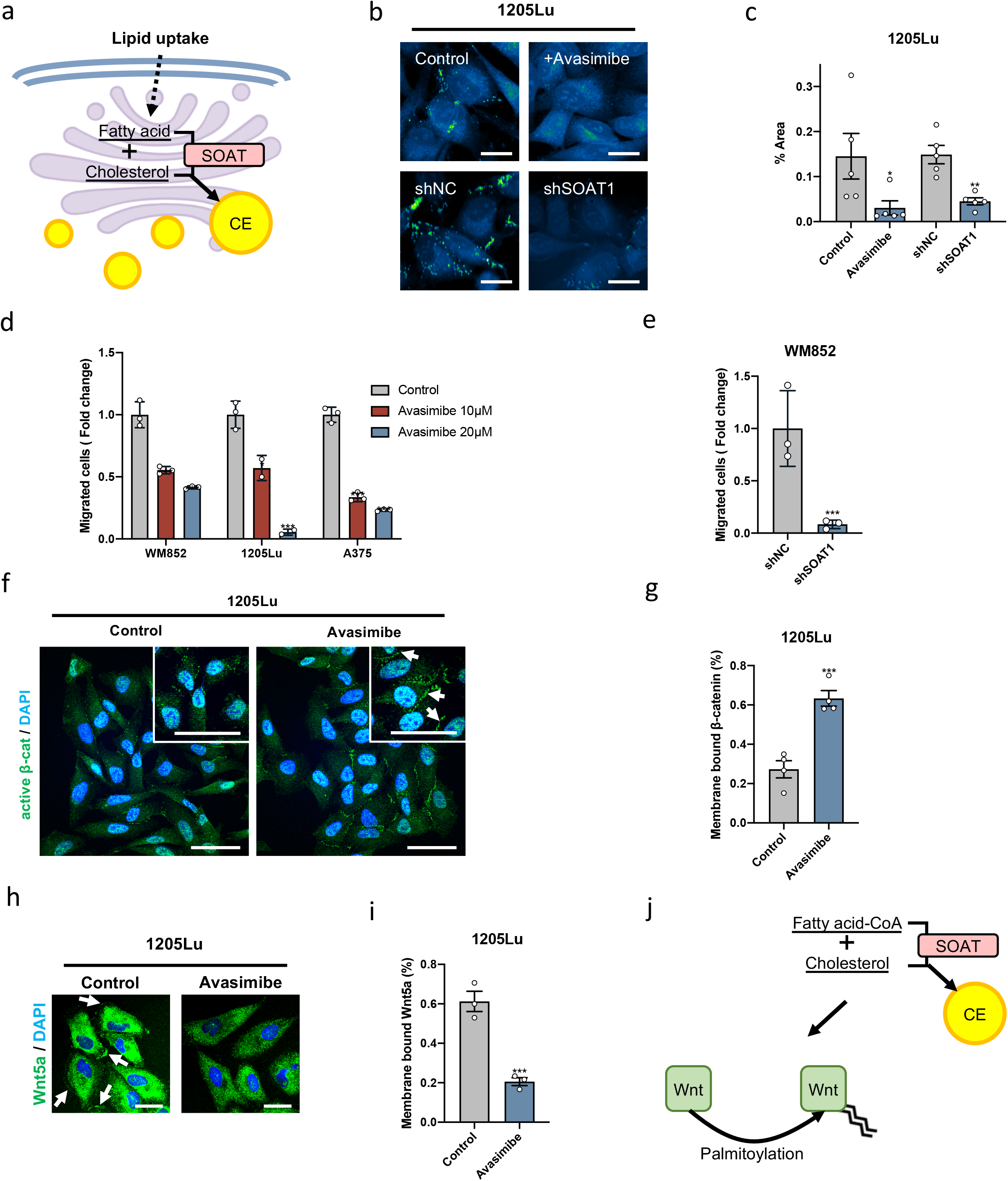
Inhibition of cholesterol esterification suppresses melanoma migration via regulation of Wnt/β-catenin pathway. (a) A schematic of cholesterol esterification process mediated by SOAT. (b) Top: Representative SRS images in the C-H region (2899 cm^−1^) of 1205Lu treated with DMSO as control and avasimibe (10 μM, 2 days). Bottom: Representative SRS images in the C-H region (2899 cm^−1^) of 1205Lu expressing negative control shRNA (shNC) and SOAT1 shRNA (shSOAT1). Scale bars: 10 μm. (c) Quantification of percent area of LDs in 1205Lu shown in (b). (d) Quantification of migrated melanoma cells treated with DMSO as control, 10 μM avasimibe and 20 μM avasimibe for 2 days. (e) Quantification of migrated WM852 expressing shNC and shSOAT1. (f) Fluorescence images of immunostaining active β-catenin in 1205Lu treated with DMSO as control and avasimibe (10 μM, 2 days). Arrows indicate membrane sequestered β-catenin. Scale bars, 50 μm. (g) Quantification of membrane bound β-catenin in 1205Lu as shown in (f). (h) Fluorescence images of immunostaining Wnt5a in 1205Lu treated with DMSO as control and avasimibe (10 μM, 2 days). Arrows indicate membrane bound Wnt5a. Scale bars, 25 μm. (i) Quantification of membrane bound Wnt5a in 1205Lu as shown in (h). (j) Proposed molecular mechanism of cholesterol esterification modulating Wnt palmitoylation. Data represent mean ± SD. *: p < 0.05, **: p < 0.01, ***: p < 0.001.

Given that inhibiting cholesterol esterification reduces the whole lipid accumulation in melanoma, we hypothesize that depleting CE reduces substrate availability for lipid modification of signaling proteins. Although cholesterol itself can modify signaling molecules such as the hedgehog protein ^37^, we did not observe inactivation of the hedgehog pathway when cholesterol esterification is inhibited (**Supplementary Fig. 14**). Instead, Wnt protein is another important signaling protein that requires lipid modification to be secreted and bound to the plasma membrane^38^. Indeed, when cholesterol esterification is inhibited by avasimibe, β-catenin is sequestered on the plasma membrane (**Fig. 6f,g**), indicating inactivation of the Wnt/β-catenin pathway. Furthermore, the amount of membrane-bound Wnt5a reduced significantly by 3-fold when cholesterol esterification is inhibited by avasimibe (**Fig. 6h,i**). A similar Wnt5a distribution was found in melanoma cells cultured with de-lipidized serum (**Supplementary Fig. 15**), which limits fatty acid uptake. Together, these results indicate that cholesterol metabolism plays an important role in melanoma metastasis, possibly mediated by a change of fatty acid availability for protein modification (**Fig. 6j**).

## Discussion

Altered lipid metabolism is being increasingly recognized in aggressive cancers ^39^. Cancer cells increase their reliance on *de novo* biosynthesis and exogenous fatty acid uptake. These lipid metabolic features provide biomolecules to sustain rapid proliferation, and at the same time, serve as an energy source during metabolic stress ^32,40^. Among the many aspects of lipid metabolism, fatty acid uptake is shown to be elevated in cancers with high metastatic potential ^41,42^. From this point of view, the metabolic microenvironment of tumors, such as adjacent adipocytes, has been shown to supply lipids and fatty acids to support cancer progression and promote metastasis ^17,43,44^. In addition to lipid degradation to promote cancer survival ^45^, fatty acids provide energy through fatty acid oxidation to enhance metastasis ^46,47^. One particular lipid species that actively contributes to cancer progression is cholesterol ^48^. Increased cholesterol biosynthesis and uptake is elevated in aggressive cancers ^49^, and the mevalonate pathway, part of cholesterol synthesis, interacts closely with oncogenic pathways ^50,51^. Accumulation of CE in LDs is found in multiple types of cancer ^52–56^, which serves as a reservoir of excess cholesterol to maintain cholesterol synthesis and uptake. These studies highlight importance of lipid metabolism during cancer progression.

Despite various efforts to understand lipid metabolic reprogramming in cancer, the alterations in composition, intracellular distribution, and dynamics of lipids are still understudied. Lipids are highly dynamic and complicated species with functional roles dependent on their distribution. For example, lipid composition changes the physical properties of the membrane, such as fluidity, which is important for cellular functions ^57,58^, protein localization and activity ^59^, cell adhesion, and migration ^35,36,60^. LDs are specialized storage organelles receiving attention ^61^ because of their multiple functions such as serving as an energy source ^62^, supporting membrane biosynthesis ^63,64^, protecting cells from lipotoxicity ^65,66^ and oxidative stress ^67^, maintaining ER homeostasis ^68,69^, and interactions with other organelles ^70^. Analyzing the composition of LDs and their spatio-temporal dynamics in cancer may suggest connections between oncogenic events and metabolic reprogramming for the development of new therapeutic targets.

High-speed, high-resolution chemical imaging enabled by coherent Raman scattering microscopy ^19,20,71^ offers an effective tool to understand the altered metabolism inside cancer cells. By SRS imaging and Raman spectroscopic analysis of single LDs in human prostate cancer patient tissues and cells, accumulation of CE in aggressive cancer was discovered, and cholesterol esterification was demonstrated as a therapeutic target of metastatic prostate cancer ^52^. Using single-cell hyperspectral SRS imaging of ovarian cancer, unsaturated lipid was identified as the metabolic signature of cancer stem cells, and inhibition of lipid desaturation can effectively eliminate cancer stem cells ^72^. Enabled by the high-speed spectral acquisition by multiplexed SRS imaging scheme, the high-content analysis using SRS imaging cytometry unveiled a metabolic response to stress in cancer cells, indicating LD-rich protrusions with high lipolysis and fatty acid oxidation activities to support cancer survival under starvation and chemotherapy ^73^. Collectively, these studies demonstrate the capability of coherent Raman scattering microscopy in understanding metabolic alterations in cancer with sub-cellular resolution and temporal dynamics.

In this study, by multimodal SRS and pump-probe imaging, metabolic reprogramming in melanoma is identified, resulting in a switch from pigmentation to LD accumulation in metastatic melanoma. Raman spectral analysis of individual LDs further reveals unsaturated fatty acids and CE. Fatty acid uptake is found to play a major role in LD accumulation and the promotion of cell migration. A human-specific fatty acid, sapienate, is found at a higher level in LD-rich metastatic melanoma and is shown to promote melanoma migration and metastasis through regulation of membrane fluidity. CE is shown to be equally important for melanoma migration. Inhibition of cholesterol esterification reduces LD accumulation and suppresses cell migration by inactivation of the Wnt/β-catenin pathway.

An intriguing question is why metastatic melanoma favors fatty acid uptake over *de novo* lipogenesis. One possible reason is that melanoma is developed in a lipid-rich local environment^17,43^. Up-regulation of several fatty acid transporter genes, such as FABP4 ^42,43^ and CD36 ^41,74,75^, has been reported in metastatic cancer. It was found that stress such as hypoxia up-regulates FABP4 expression ^42^. Following enhanced fatty acid uptake, fatty acid oxidation is found to play an important function in cancers. For example, fatty acid oxidation helps cancer cells to survive under metabolic stress ^40,73^. In melanoma, fatty acid oxidation-dependent drug resistance has been reported ^76^. Inhibition of fatty acid oxidation suppresses metastasis to local lymph nodes ^46^, indicating the importance of fatty acids in promoting local metastasis in melanoma. Intriguingly, blood-born metastasis to the lungs was not suppressed by fatty acid oxidation inhibition ^46^, suggesting other functions of fatty acids exist that promote melanoma metastasis beyond local lymph nodes. In this study, we identified another important function of fatty acids in the cell, which is through the regulation of membrane fluidity and post-translational modifications of proteins.

There are two major fatty acid monodesaturation pathways: SCD-mediated, generating palmitoleate, and FADS2-mediated, generating sapienate. SCD is essential for cancer cell survival, and its anti-cancer potential has been reported in multiple cancers ^77^. However, not all cancer cells are responsive to SCD inhibition ^39^, and a recent study demonstrated FADS2 as an alternative desaturation pathway in SCD-independent cancer cells ^31^. Unlike other fatty acids, sapienate is only found in humans and is a major component of human sebum ^78^. Intriguingly, a recent study indicated that SCD expression is positively regulated by MITF ^79^. Therefore, one potential mechanism of increased dependency on FADS2-mediated fatty acid desaturation in MITF^low^/AXL^high^ LD-rich melanoma cell lines could be via MITF. Importantly, as the high migration and invasion capacities presented in the melanoma can be suppressed by FADS2 inhibition *in vitro* and *in vivo*, inhibition of FADS2-mediated fatty acid desaturation offers a therapeutic opportunity to treat metastatic melanoma.

In addition to serving as a Δ6 desaturase to generate sapienate from palmitate, FADS2 mediates desaturation to generate polyunsaturated fatty acids ^80^. In metastatic melanoma, this sapienate synthetic enzyme is a major contributor promoting cell migration. This function is supported by the capacity of sapienate to rescue cell migration capacity in the de-lipidized serum condition. Further, the higher amount of sapienate found in LD-rich melanoma compared to LD-poor melanoma suggests elevated sapienate synthesis in metastatic melanoma. Mechanistically, unsaturated lipids on the plasma membrane are known to promote cell mobility ^57,58^. Importantly, FADS2 displays a plasma membrane expression pattern, which is not commonly found in fatty acid desaturases. In line with this observation, the membrane fluidity is higher in LD-rich melanoma compared to LD-poor melanoma, and supplementation of sapienate increases membrane fluidity. These results collectively indicate an essential function of sapienate in cancer migration.

In parallel to the reprogramming of fatty acid metabolism in metastatic melanoma, cholesterol metabolism also plays an essential role in promoting cell migration. A significant amount of CE in LDs of metastatic melanoma suggests a rewiring of cholesterol metabolism. With the increasing number of reports regarding the up-regulation of SOAT-mediated cholesterol esterification in cancers ^52–56^, it is essential to understand the physiological meaning and function of CE accumulation in metastatic disease. Our results demonstrate that SOAT inhibition effectively suppresses melanoma migration. Based on the reduced LD accumulation observed in SOAT-inhibited melanoma, it is likely that cholesterol esterification is connected to fatty acid uptake, storage and utilization. Fatty acid is known to serve as a substrate for protein modification such as palmitoylation to regulate protein localization and activities. For example, Wnt is one of the proteins that requires palmitoylation to be transported to the plasma membrane for secretion ^38^. In this study, we demonstrate that SOAT inhibition could inactivate Wnt5a in metastatic melanoma via reducing membrane localization of Wnt5a. As Wnt5a is a robust marker of aggressive and metastatic melanoma ^81^, our finding of modulation of Wnt5a modification by reprogrammed lipid metabolism provides an essential mechanism of linking metabolism with signaling pathways.

Finally, the metabolic reprogramming profile identified in this study serves as a signature for metastatic melanoma diagnosis. Without specific molecular markers, clinical diagnosis of invasive melanoma is often challenging. It was found that more eumelanin than pheomelanin is present in malignant melanoma ^6^. Using pump-probe imaging, the distribution of these two types of melanin was studied in human tissues, which provide a label-free imaging approach for melanoma diagnosis ^8^. More recently, chemical analysis of melanin with its spatial distribution using pump-probe microscopy further demonstrated the potential of predicting the metastatic capability of melanoma ^9^. Despite these advances, loss of MITF found in melanomas with an invasive phenotype ^10^ suggests loss of pigmentation, indicating a need to identify new molecular markers for detecting aggressive and invasive melanoma. In this study, by integration of label-free SRS imaging of lipids and pump-probe imaging of pigments, we identified a metabolic switch from pigmentation to lipid accumulation in metastatic melanoma. Such a multimodal imaging platform can potentially be used to diagnose primary melanoma and metastatic lesions in human tissue samples. Furthermore, with the continued development of a handheld SRS imaging device ^82^, *in vivo* SRS and pump-probe imaging promises a non-invasive approach for clinical evaluation of melanoma.

## Materials and Methods

### Cell lines and cell culture

Human melanoma cell lines (WM35, WM902B, WM983A, WM983B, 1205Lu, WM793, A375) harboring the BRAF V600E mutation and human melanoma cell line (WM852) with wildtype BRAF were obtained from Dr. Meenard Herlyn (The Wistar Institute). Cells were grown in Dulbecco’s Modified Eagle’s Medium (DMEM, Invitrogen) media supplemented with fetal bovine serum (FBS) (10%), L-glutamine (2mM), penicillin (1%), and streptomycin (1%). Human primary melanocytes (Life Technologies) were grown in Medium 254 (Gibco) with human melanocyte growth supplements (Gibco). Cell lines were incubated at 37 °C in 5% CO_2_.

### Chemicals

FBS was purchased from Life Technologies. De-lipidized serum was purchased from Gemini Bio. CAY10566, BMS309403, Sulfosuccinimidyl Oleate, lipofermata, and etomoxir were purchased from Cayman Chemical. SC26196 was purchased from Santa Cruz. Avasimibe was purchased from Selleckchem. Glucose-d_7_, palmitic acid-d_31_, and oleic acid-d_34_ were purchased from Cambridge Isotope Laboratories. Palmitate, palmitoleate, oleate, cholesteryl oleate, and glyceryl trioleate were purchased from Sigma-Aldrich. Sapienate was purchased from Matreya. Octadecenoate (8(Z)-Octadecenoic acid) was purchased from Larodan.

### RT-qPCR

RNA was isolated following the RNeasy Plus Mini Kit protocol (Qiagen). RNA concentration was quantified using the NanoDrop™ (Thermo Fisher Scientific), and 1 μg RNA was reverse transcribed and amplified using the Superscript™ III First-Strand Synthesis System (Thermo Fisher Scientific). Resultant cDNA (1μL) was added to primer working solutions (IQ™ SYBER^®^ Green Supermix, UltraPure™ Distilled Water, and forward and reverse primer mix) for each well in the 96-well qPCR plate (**Table 1**). Amplification and quantification were performed using the StepOnePlus™ Real-Time PCR System (Thermo Fisher Scientific). All reactions were performed in triplicate, using GAPDH as an internal control. Results were quantified as Ct values, which represent the threshold cycle of PCR at which the amplified product is first detected, and expressed as the ratio of target/control (relative gene expression) using the 2^−ΔΔCt^ method.

**Table 1.**
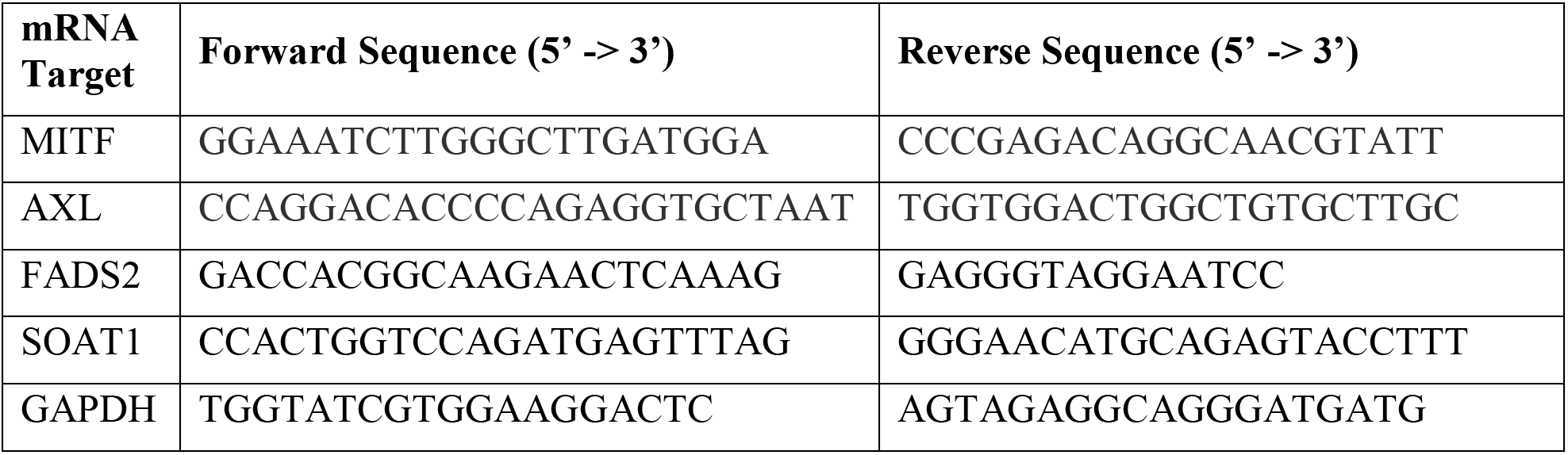
Primer sequences used for RT-qPCR.

### Time-domain multimodal SRS/pump-probe imaging

SRS/pump-probe imaging was performed on a femtosecond SRS microscope. An ultrafast laser system with dual output at 80 MHz (InSight DeepSee, Sepctra-Physics) provided pump and Stokes beams. In SRS imaging scheme, 802 nm and 1045 nm beams serve as pump and Stokes, respectively, to be resonant with the C-H stretching vibration at 2899 cm^−1^. For pump-probe imaging scheme, 802 nm and 1045 nm beams serve as probe and pump beams, respectively. For off-resonance imaging of the C-H stretching vibration, the pump beam was tuned to 845 nm, which corresponds to 2265 cm^−1^. In this imaging scheme, pump-probe signal, generated by 1045 nm pump and 845 nm probe, becomes dominant because no endogenous Raman active biomolecules are presented in the samples. 1045 nm beam was modulated by an acousto-optic modulator (AOM, 1205-C, Isomet) at 2.2 MHz. Both beams were linearly polarized. A motorized translation stage was employed to scan the temporal delay between the two beams. Two beams were then sent into a home-built laser-scanning microscope. A 60x water immersion objective lens (NA = 1.2, UPlanApo/IR, Olympus) was used to focus the light into the sample, and an oil condenser (NA = 1.4, U-AAC, Olympus) was used to collect the signal. The stimulated Raman loss and pump-probe signals were detected by a photodiode, which was extracted with a digital lock-in amplifier (Zurich Instrument). The power of the tunable beam (802 nm and 845 nm) and the power of 1045 nm beam at the specimen were maintained at ~10 mW and 5 to 25 mW, respectively. The images were acquired at 10 μs pixel dwell time. No cell or tissue damage was observed during the imaging procedure.

### Phasor analysis for LD and pigment quantification

LDs and pigments were analyzed by phasor analysis using ImageJ plugin ^27,83^. The segmentation of LDs and pigments in the phasor space is shown in the **Supplementary Fig. 4**. After the segmentation, “Threshold” function was used to select droplets in the cells. “Analyze Particles” function was then used to quantify the area fractions of droplets in the whole image area, then normalized to the cell number counted from the same image.

### *In silico* analysis of melanoma using TCGA datasets

The transcriptome dataset of melanoma tissues (TCGA-SKCM) obtained from TCGA was downloaded in Apr 2019 using the UCSC Xena browser ^84^. The patients were assigned to MITF^high^/AXL^low^ and MITF^low^/AXL^high^ groups based on the median expression level. Differentially expressed genes and Gene Set Enrichment Analysis analyses were generated using NetworkAnalyst 3.0 ^85^. Further enrichment analyses were performed in the included genes (FC > 1.2, adjusted p < 0.05) using Metascape ^86^.

### Human tissue sample preparation

Deidentified frozen specimens of human primary and metastatic melanoma tissues were purchased from IU Simon Comprehensive Cancer Center (Lafayette, IN). The tissue samples were sliced using a cryostat at 10 to 20 μm thickness. The time-domain multimodal SRS/pump-probe imaging was performed on these tissue slices without any processing or labeling.

### Confocal Raman spectral measurement

Raman spectral analysis from individual LDs was performed using a commercial confocal Raman microscope (LabRAM HR Evolution, Horiba) at room temperature. A 15 mW (after the objective), 532-nm diode laser was used to excite the sample through a 40× water immersion objective (Apo LWD, 1.15 N.A., Nikon). The total data acquisition time was 60 s using the LabSpec 6 software. For each group, at least 10 spectra from individual LDs in different locations or cells were obtained. To analyze the spectrum, the background was removed manually based on the glass background profile, and peak intensity was measured using OriginPro. The calibration curve for CE percentage in LD was generated by measuring CE/triacylglyceride emulsion at various percentages and linearly correlate it with the 704 cm^−1^ peak (cholesterol rings) normalized with the 1445 cm^−1^ peak (CH_2_ bending vibration). Unsaturation degree of LD is determined by the peak intensity at 1654 cm^−1^ (C=C stretching vibration) normalized with the 1445 cm^−1^ peak.

### Metabolite extraction and measurement by LC/MS and GC/MS

The cells were extracted using a modified Bligh-Dyer extraction ^87^. To each sample, 0.4 mL of ice cold 65% methanol was added and vortexed for 1 min. The samples were placed on ice, spiked with 1 μg of C17:0 margaric acid as internal standard, then pulse vortexed to mix. Next, 0.25 mL of chloroform was added and the sample vortexed for an additional 5 minutes. The samples were centrifuged at 13,500 rpm for 5 minutes at 4°C. The bottom layer was removed for fatty acid analysis. The samples were dried using a rotary evaporation device at room temperature for 2 hours. The samples were then derivatized for GC/MS analysis. Each sample received 0.5 mL of 14% boron trifluoride solution (BF3) (Sigma-Aldrich # B1252) and reacted for 30 minutes at 60°C. Then 0.25 mL of water and 1 mL of hexane were added. The samples were mixed, then dried with approximately 0.2g of anhydrous sodium sulfate. The hexane layer was then collected and dried using a stream of nitrogen. The final derivatized sample was reconstituted in 0.1 mL of hexane for GC/MS analysis.

A Thermo Fisher Triplus RSH auto sampler and Trace 1310 gas chromatography (GC) system coupled to a Thermo Fisher TSQ 8000 mass spectrometer (MS) was used to analyze FAME composition in each sample (Thermo Fisher Scientific, Waltham, MA). An Agilent Select FAME GC column (50 m x 0.25 mm, film thickness 0.2 um) was used for the analysis (Agilent Technologies, Santa Clara, CA). The GC carrier gas was helium with a linear flow rate of 1.0 ml/min. The programmed GC temperature gradient was as follows: time 0 minutes, 80°C, ramped to 175°C at a rate of 13°C/min with a 5-minute hold, then ramped to 245°C at a rate of 4°C/min with a 2-minute hold. The total run time was 38.3 minutes. The GC inlet was set to 250°C and samples were injected in split-less mode. The MS transfer line was set to 250°C and the MS ion source was set to 250°C. MS data were collected in selected ion monitoring (SIM) mode according to **Table 2**. All data were analyzed with Thermo Fisher Chromeleon (Version 7.2.9) software. A standard mixture of 37 FAME (Supelco, Sigma-Aldrich), sapienate (Larodan), and palmitoleate (Sigma-Aldrich) were used to confirm spectra and column retention times.

**Table 2.**
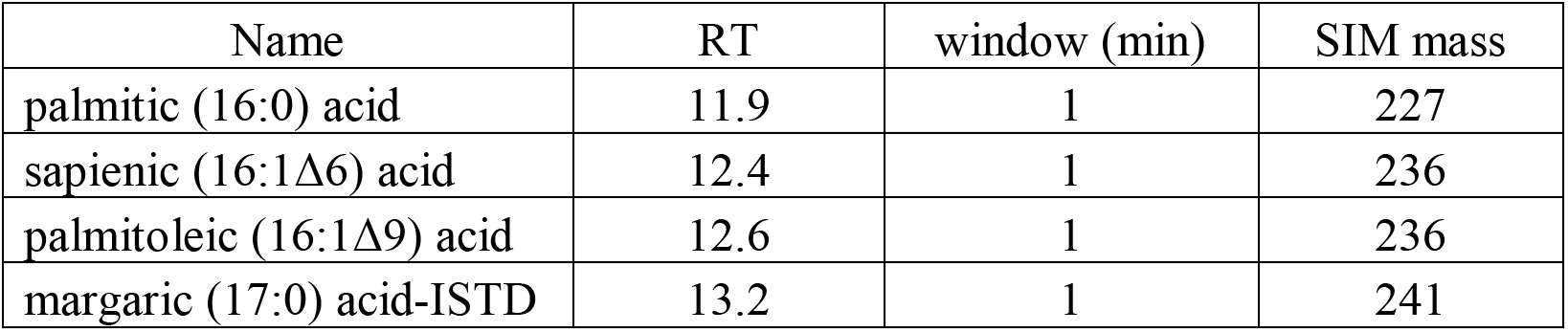
SIM table for mass spec analysis.

### Isotope metabolite labeling experiments

For tracking *de novo* lipogenesis, cells are cultured with glucose-d_7_ (4.68 g/L) in the glucose-free complete medium for 48 hours. For tracking fatty acid uptake, 100 μM of palmitic acid-d_31_ or oleic acid-d_34_ was supplemented in the complete medium and cultured for 6 hours. The pump laser was tuned to 855 nm to perform SRS imaging in the C-D region at 2127 cm^−1^. Pump and Stokes powers at the specimen were maintained at ~10 mW and 50 mW, respectively. To quantify the SRS intensity at C-D region from LDs, “Threshold” function in ImageJ was used to select LDs in the cells. Then the total intensity of LDs was obtained after particle analysis and normalized by the number of cells in the corresponding image.

### Migration/invasion assay

0.2 million cells were seeded in the upper chamber of a 24-well plate insert (Corning™ Transwell™ 8 μm Permeable Polycarbonate Membrane Inserts) in serum-free media. For invasion assay, the insert was pre-coated with Matrigel (BD Biosciences). DMEM media supplemented with 20% FBS and 50 ng/mL EGF was added to the bottom chamber of the insert. For the treatment group, inhibitors or fatty acids was added to both the upper and bottom chambers. After 4.5 to 8 hours of migration, cells were fixed with either 10% formalin or 70% ethanol and non-migrated cells removed with a cotton swab. Migrated cells were stained with 50 μg/mL propidium iodide (Invitrogen) for 30 minutes at room temperature and then washed with phosphate buffered saline 5 to 6 times. 8 to 10 random images were taken for each well using an FV3000 confocal microscope or a Nikon Eclipse E400 microscope. The number of migrated/invaded cells was quantified using ImageJ. Migrated cell count was normalized to the control group and represented as fold change from control unless indicated otherwise.

### Extracellular flux analysis

Etomoxir was dissolved in sterile water to a final stock concentration of 2 mM. The Etomoxir stock solution was aliquoted and stored at −20°C. Fresh Etomoxir aliquots were used for each treatment, and treatments were refreshed every 2-3 days. Cells (15,000 cells/well) were plated on an XF96 polystyrene cell culture microplate (Seahorse®, Agilent). The following day, cells were washed and incubated in XF DMEM pH 7.4 assay medium supplemented with 10 mM glucose, 1 mM pyruvate, and 2 mM glutamine (Seahorse®, Agilent) in a CO_2_ free incubator for 1 hour. The plate was inserted into the Seahorse Xfe96 Extracellular Flux Analyzer, and injections of etomoxir (4 μM final concentration), Oligomycin (1.5 μM final concentration), and Antimycin A (1 μM final concentration) were sequentially added to assess the role of mitochondrial fatty acid oxidation on ATP generation. Simultaneous oxygen consumption rate (OCR) and extracellular acidification rate (ECAR) measurements were obtained every 3 minutes; a minimum of three measurements was acquired after each injection. OCR (pmol O2/min) and ECAR (mpH/min) values were normalized to DNA content (ng/μL) measured using the PicoGreen™ Assay. The Seahorse® Wave V2.6 software was used for data analysis.

### Knockdown strategies

1205Lu cells were transfected with FADS2 targeting shRNA lentiviral particles (Applied Biological Materials, shFADS2 #1: CCGGCCACGGCAAGAACTCAAAGATCTCGAGATCTTTGAGTTCTTGCCGTGGTTTTTG, shFADS2 #2: CCGGCCAC CTGTCT-GTCTACAGAAACTCGAGTTTCTGTAGACAGACAGGTGGTTTTTG) following the protocol provided by the manufacturer. Scrambled shRNA lentiviral particles (shNC: GTCTCCACGCGCAGTACATTT) were used as a control. Stably transfected cells were selected with 1 μg/mL puromycin. 1205Lu and WM852 cells were transfected with SOAT1 targeting shRNA lentiviral particles (Santa Cruz, sc-29624-V) following the protocol provided by the manufacturer. Stably transfected cells were selected with 1 μg/mL puromycin.

### Mouse model and fluorescence imaging of tissue sections

All animal procedures were approved by Boston University IACUC. The mouse tail-vein injection model was used to study the development of metastatic cancer. 6-week old male homozygous nude (Foxn1^nu^/Foxn1^nu^) mice obtained from the Jackson Laboratory were used. 1205Lu melanoma cells stabling expressing GFP (Applied Biological Materials) were transfected with lentivirus carrying negative control shRNA (1205Lu-shNC) or FADS2 targeting shRNA (1205Lu-shFADS2 #2). The cells were then prepared in sterile PBS in a completely monocellular suspension without clumps at 1 × 10^6^/mL concentration. 100 μL of cell suspension was injected slowly via the lateral tail vein of the anesthetized mouse, after which the bleeding was stopped by applying pressure to the puncture site with a dry piece of gauze. The lung tissues were collected 46 days after tumor inoculation, and sliced using a cryostat at 10 to 20 μm thickness. Then the tissues were mount with the antifade mounting medium with DAPI (Vector Laboratories). The tissues were imaged using an Olympus VS120 Automated Slide Scanning Microscope.

### Immunofluorescence staining and imaging

Cells were fixed with 10% formalin for 15 min at room temperature. Cells were then incubated with anti-adipophilin (Millipore Sigma, 393A-1, 1:100), anti-FADS2 (Proteintech, 28034-1-AP, 1:200), anti-Gli1 (Abcam, ab49314, 1:200), anti-active β-catenin (Millipore Sigma, 05-665, 1:200) or anti-Wnt5a (Santa Cruz, sc-365370, 1:200). After incubation with the secondary antibody containing Alexa-Fluor 488, cells mount using the antifade mounting medium with DAPI (Vector Laboratories). Then the cells were imaged under an FV3000 confocal microscope with a 60x oil objective. Fluorescence signal from the antibody was imaged with 488 nm excitation and 500 - 600 nm emission. DAPI signal was imaged with 405 nm excitation and 430 – 479 nm emission.

### Membrane fluidity assay

Membrane fluidity was measured using MarkerGene™ Membrane Fluidity Kit (Abcam) following the protocol provided by the manufacturer. Briefly, cells were grown on 96-well plates. After 24 to 48 hours, the cells were washed with PBS and incubated with 10 μM of pyrenedecanoic acid (PDA) in Perfusion Buffer containing 0.08% pluronic F127 for 20 minutes at 25°C in the dark. Then the unincorporated PDA is removed by washing the cells with serum-free media twice. The incorporated PDA was measured by reading fluorescence signals at both 400 (monomer) and 460 nm (excimer) with excitation at 360 nm using SpectraMax i3x Microplate Detection Platform (Molecular Devices). The membrane fluidity was represented as a ratio between excimer fluorescence and monomer fluorescence, in which a higher ratio indicates higher membrane fluidity.

### Confocal imaging of membrane fluidity

For imaging membrane fluidity, cells were grown on glass coverslips for 48 hours. Cells were stained with di-4-ANEPPDHQ (Thermo Fisher Scientific) following the protocol provided by the manufacturer. Briefly, 5 mM of Di-4-ANNEPPDHQ stock solution in DMSO was diluted in serum-free DMEM (final concentration: 5 μM). Cells were incubated with Di-4-ANNEPPDHQ-containing DMEM for 30 minutes at 37°C in the dark. Then cells were washed with DMEM and imaged using an FV3000 confocal microscope with excitation at 488 nm and emission at both 530 – 590 (ordered lipid) and 590 – 650 nm (disordered lipid). The images were analyzed using ImageJ to generate ratiometric results of ordered-to-disordered lipids.

### Statistical analysis

Statistical analysis was performed using Origin Pro or Prism. For two-sample comparisons, data was first tested for normality. If the data is detectably non-Gaussian, a nonparametric Mann-Whitney U test was performed. Otherwise, a one-tailed student’s t-test was performed. Probability of the null hypothesis P < 0.05 was judged to be statistically significant.

### Reporting Summary

Further information on research design is available in the Nature Research Reporting Summary linked to this article.

## Data availability

Relevant data analyzed during this study are included in this article and its Supplementary Information files. All additional data that support the findings are available from the corresponding author upon request.

## Acknowledgement

This research is supported by NIH grant R33 CA223581 to JXC. The authors thank the IUSCC Cancer Center at Indiana University School of Medicine funded by the IU Simon Cancer Center Support Grant P30 CA082709, for the use of the Tissue Procurement and Distribution Core, which provided Frozen Tissue Sample service. The research reported in this publication was supported by the Boston University Micro and Nano Imaging Facility and the Office of the Director, National Institutes of Health of the National Institutes of Health under award Number S10OD024993. The content is solely the responsibility of the authors and does not necessarily represent the official views of the National Institute of Health. The Seahorse XFe96 Extracellular Flux experiments were supported by the Boston University Analytical Instrumentation Core. The authors acknowledge the use of the facilities of the Bindley Bioscience Center, a core facility of the NIH-funded Indiana Clinical and Translational Sciences Institute. The authors also acknowledge the Animal Science Center for providing technical support for animal study reported within this paper. The authors thank Kai-Chih Huang, Peng Lin, Hongli Ni, and Haonan Lin for their helps on SRS imaging and pump-probe imaging; thank Junjie Li and Yuying Tan for their helpful discussions; thank Amber S. Jannasch and Bruce Cooper from Purdue Metabolomics Facility for their helps on mass spectrometry measurements.

## Author Contributions

HJL, JXC, RA co-designed the experiments. HJL and ZC performed the experiments and data analysis. MC and JGC performed experiments and data analysis related to fatty acid oxidation. MW and MC performed measurements for MITF/AXL expressions on melanoma. HJL, JXC, ZC co-wrote the manuscript. All authors have read and commented on the manuscript.

