## Supplementary Figure for "FADS2-mediated fatty acid desaturation and cholesterol esterification are signatures of metabolic reprogramming during melanoma progression"

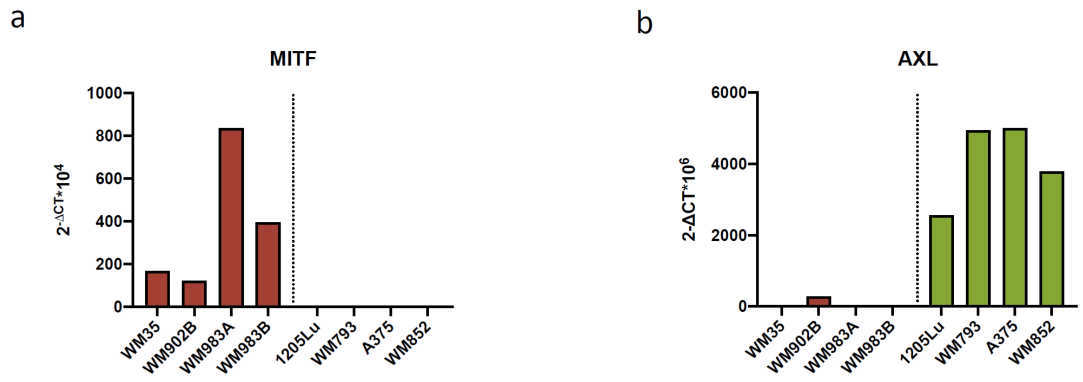


**Supplementary Figure 1. MITF and AXL expressions are inversely correlated to each other in human melanoma cell lines.** (a) MITF mRNA levels in a panel of human melanoma cell lines. (b) AXL mRNA levels in the same panel of human melanoma cell lines.

**Supplementary Table 1. Summary of human melanoma cell lines used in the study.**

| Name | Clinical data | Stage | BRAF | MITF/AXL | Lipid | Pigment | Migration | Fatty acid uptake |
|---|---|---|---|---|---|---|---|---|
| WM35 (RRID:CVCL_0580) | 24 years female | RGP | V600E | $MITF^{high}$/ $AXL^{low}$ | Low | Some | Low | - |
| WM902B (RRID:CVCL_6807) | unknown male | VGP | V600E | $MITF^{high}$/ $AXL^{low}$ | Low | High | Low | Low PA-$d_{31}$ Low OA-$d_{34}$ |
| WM983A (RRID:CVCL_6808) | 54 years male | VGP | V600E | $MITF^{high}$/ $AXL^{low}$ | Low | High | Low | Low PA-$d_{31}$ |
| WM983B (RRID:CVCL_6809) | 54 years male | MET | V600E | $MITF^{high}$/ $AXL^{low}$ | Low | High | Low | Low PA-$d_{31}$ |
| WM852 (RRID:CVCL_6804) | 63 years male | MET | WT | $MITF^{low}$/ $AXL^{high}$ | High | None | High | High PA-$d_{31}$ |
| 1205Lu (RRID:CVCL_5239) | 37 years male | Xenograft MET | V600E | $MITF^{low}$/ $AXL^{high}$ | High | Low | High | High PA-$d_{31}$ High OA-$d_{34}$ |
| WM793 (RRID:CVCL_8787) | 37 years male | VGP | V600E | $MITF^{low}$/ $AXL^{high}$ | High | Low | High | High PA-$d_{31}$ |
| A-375 (RRID:CVCL_0132) | 54 years female | MET | V600E | $MITF^{low}$/ $AXL^{high}$ | High | None | High | - |

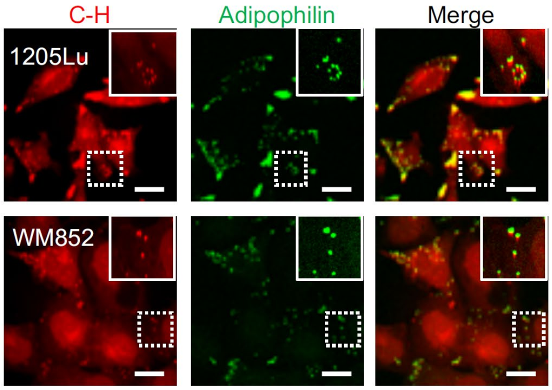


**Supplementary Figure 2. Droplets identified in MITF$^{low}$/AXL$^{high}$ melanoma cell lines are confirmed to be LDs by immunofluorescence.** Simultaneous SRS imaging in the C-H region and immunofluorescence analysis of adipophilin (green) in 1205Lu and WM852 cells. Scale bars, 10 μm.

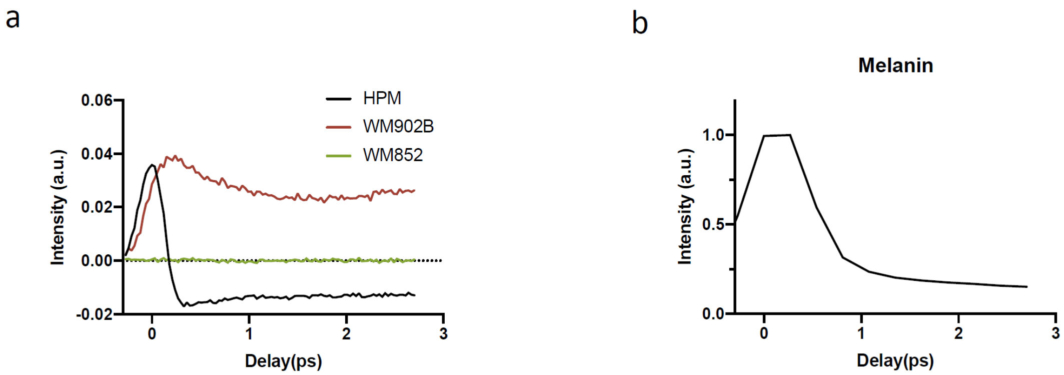


**Supplementary Figure 3. Pump-probe signals from pigments in melanocyte, melanoma, and purified melanin.** (a) Time-resolved pump-probe signals from droplets in HPM (human primary melanocyte), WM902B ($MITF^{high}/AXL^{low}$), and WM852 ($MITF^{low}/AXL^{high}$) cells. The curves were subtracted with the background generated from cross-phase modulation. No pump-probe signals were found in WM852 cells. (b) Time-resolved pump-probe signal from eumelanin drived from *Sepia Officinalis*.

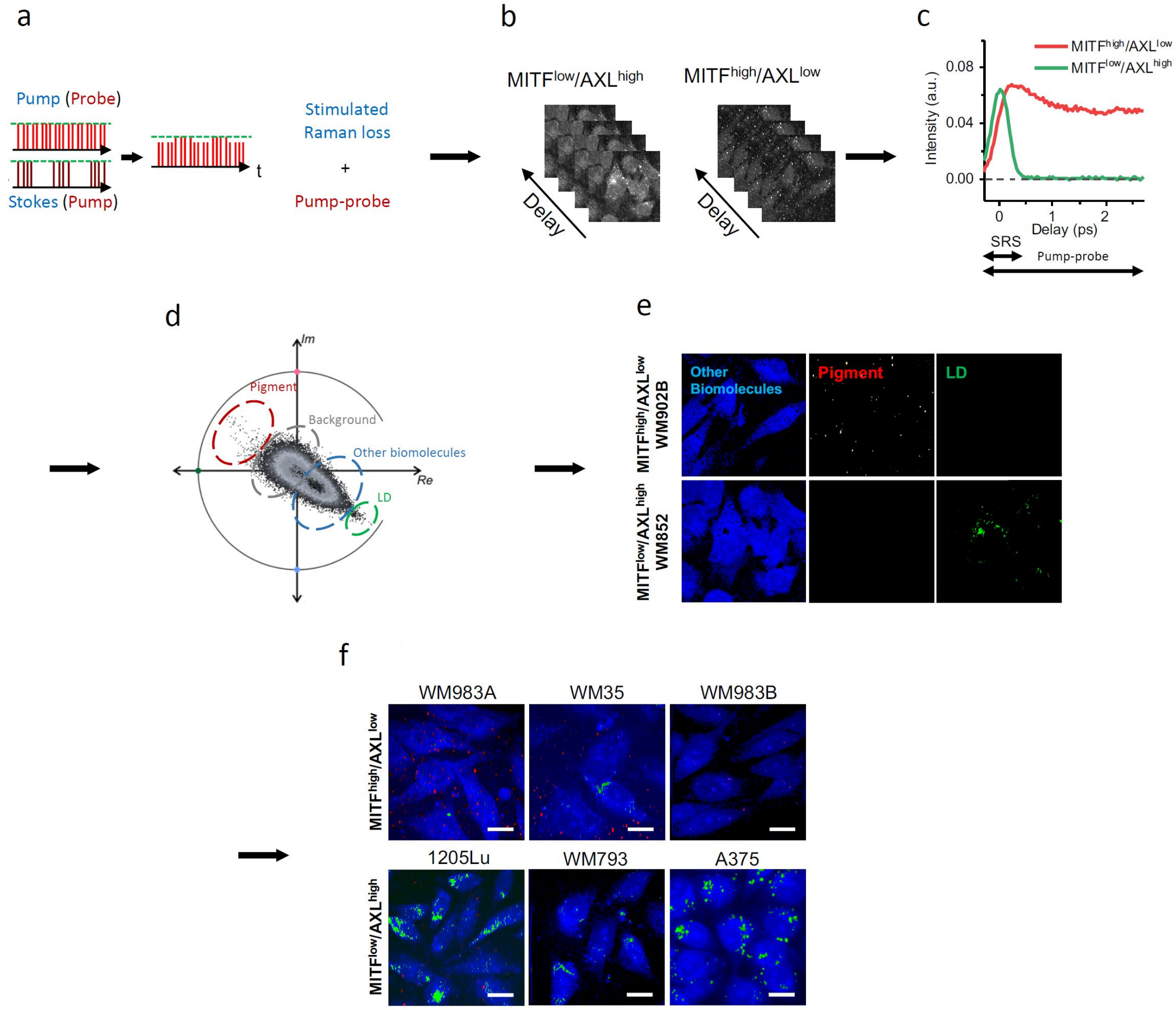


**Supplementary Figure 4. Pigments and LDs can be separated by time-domain multimodal SRS/pump-probe imaging and phasor analysis.** time-resolved measurement, and phasor analysis. (a) A schematic of SRS/pump-probe imaging. (b) A frame-by-frame time-resolved imaging was performed on MITF$^{low}$/AXL$^{high}$ and MITF$^{high}$/AXL$^{low}$ melanoma. (c) Time-resolved SRS and pump-probe signals from droplets in MITF$^{high}$/AXL$^{low}$ and MITF$^{low}$/AXL$^{high}$ melanoma. (d) A phasor plot of the time-resolved images of MITF$^{high}$/AXL$^{low}$ and MITF$^{low}$/AXL$^{high}$ melanoma. The regions representing pigments, background, lipid droplets and background are indicated. (e) Phasor output of multimodal SRS/pump-probe images. (f) Phasor output of multimodal SRS/pump-probe images from melanoma cell lines grouped based on MITF/AXL status. Scale bars, 20 µm. SRS: stimulated Raman scattering; LD: lipid droplets.

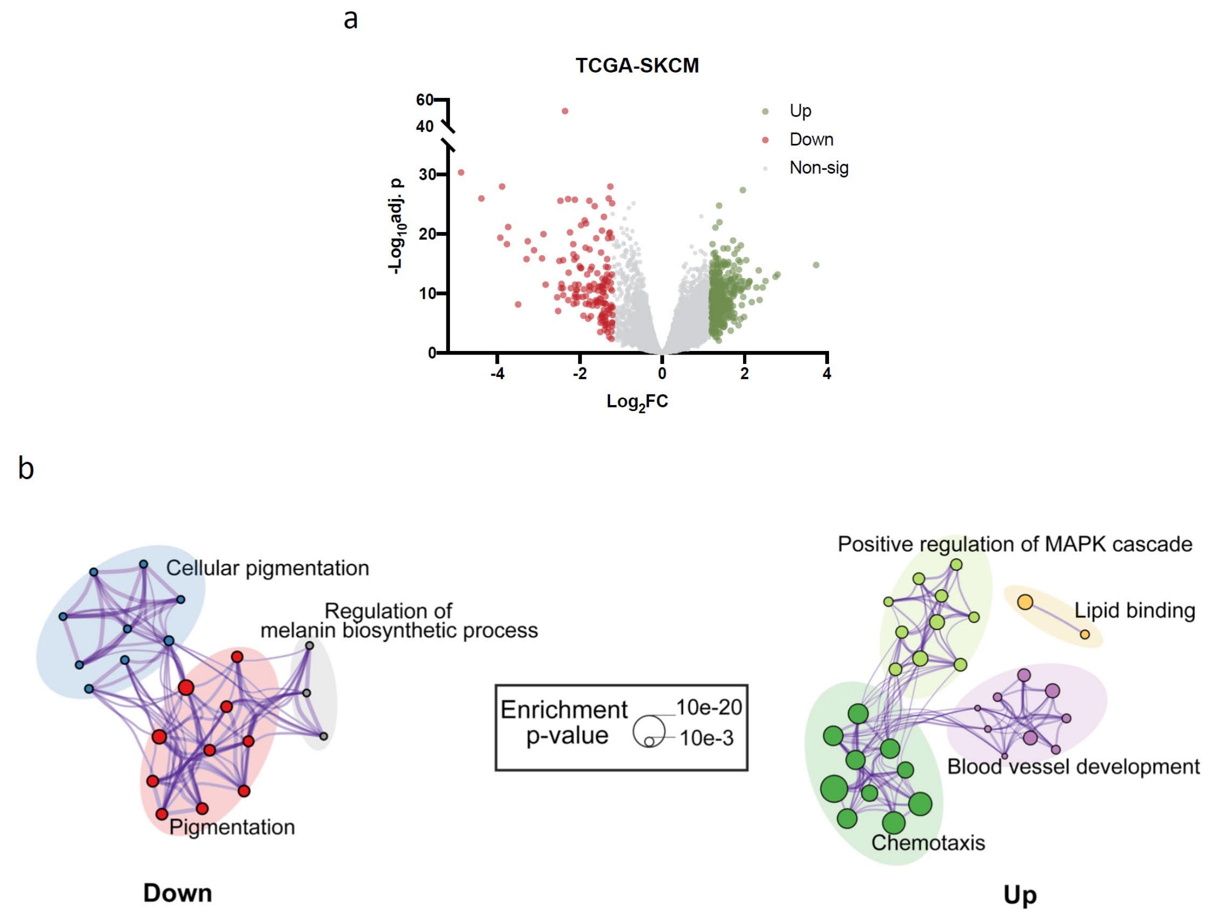


**Supplementary Figure 5.** (a) Differentially expressed genes (DEGs) between MITF$^{low}$/AXL$^{high}$ and MITF$^{high}$/AXL$^{low}$ patients in TCGA-SKCM metastatic subgroup. (b) Enrichment analysis results for the DEG between MITF$^{low}$/AXL$^{high}$ and MITF$^{high}$/AXL$^{low}$ patients. Individual gene ontology terms with similar gene members and clustered by categories indicated by node colors and labeled using a representative member. Terms with a Kappa similarity score > 0.3 are connected by edges. Node size is proportional to enrichment p-value.

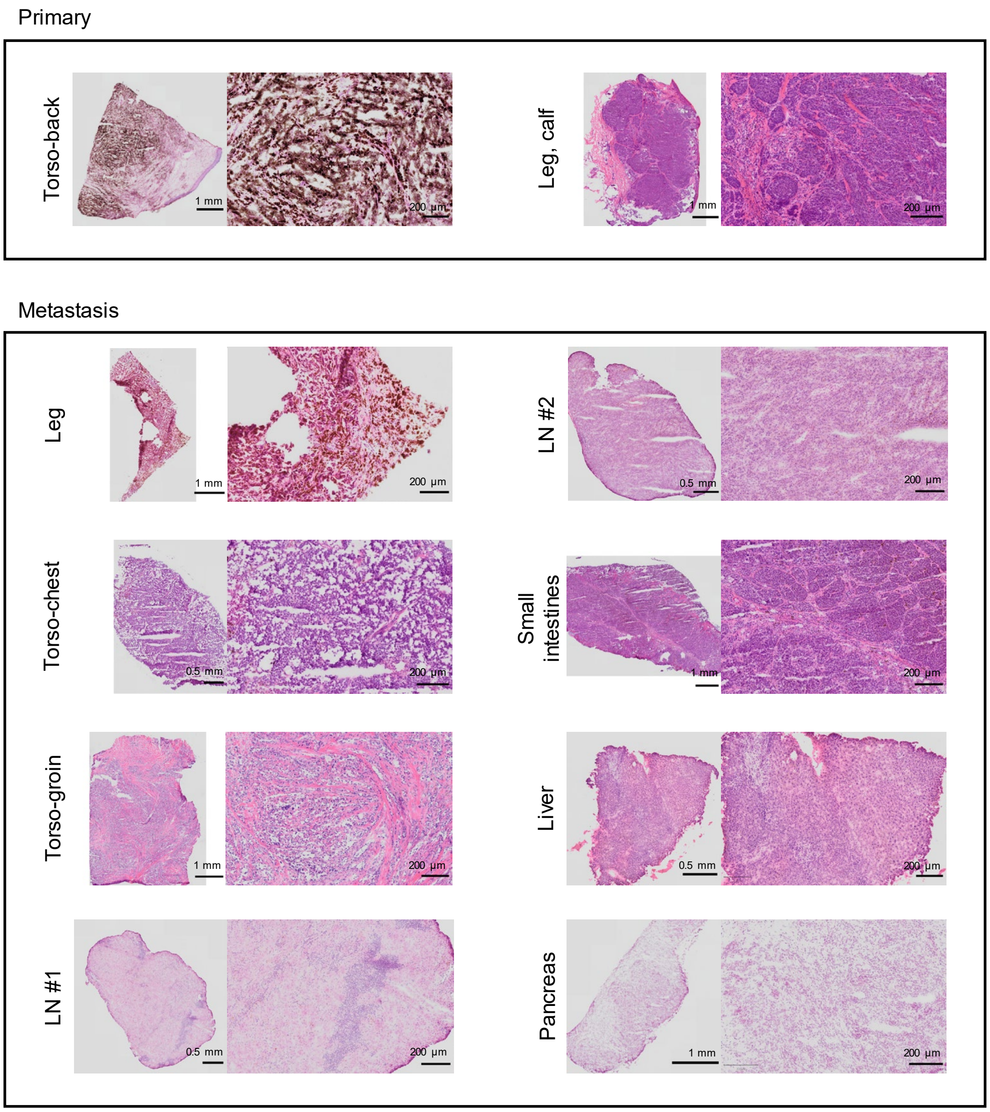


**Supplementary Figure 6.** Hematoxylin and eosin (H&E) staining of adjacent slices used for multimodal SRS imaging in **Fig. 1e**.

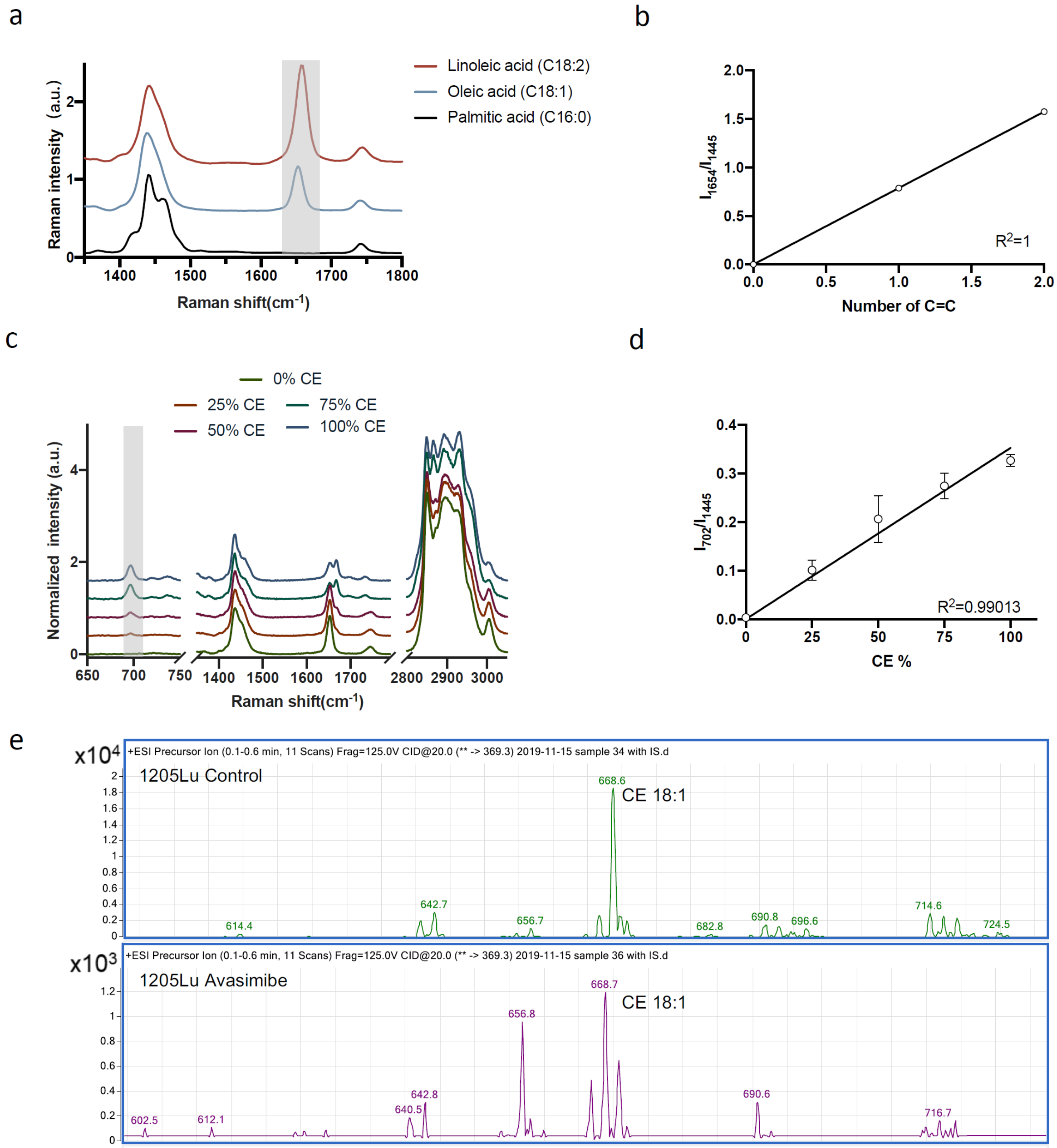


**Supplementary Figure 7. LDs in MITF$^{low}$/AXL$^{high}$ melanoma cells contain unsaturated fatty acids and CE.** (a) Representative Raman spectra of palmitic acid, oleic acid, and linoleic acid. The peak representing C=C bond is highlighted in grey. (b) Calibration curve for quantification of unsaturation degree based on the number of C=C bonds, generated by linear fitting of height ratio between the peak at 1654 cm$^{-1}$ ($I_{1654}$) and the peak at 1445 cm$^{-1}$ ($I_{1445}$). $I_{1654}/I_{1445} = 0.788 \times$ number of C=C. (c) Representative Raman spectra of CE and triacylglycerol emulsions with five different molar percentages of CE, ranging from 0% to 100%. Emulsions are mixtures of cholesteryl oleate and glyceryl trioleate. (d) Calibration curve for quantification of molar percentage of CE out of total lipid, generated by linear fitting of height ratio between the peak at 702 cm$^{-1}$ ($I_{702}$) and the peak at 1445 cm$^{-1}$ ($I_{1445}$). $I_{702}/I_{1445} = 0.00353 \times$ CE percentage. Data adopted from ref [1]. (e) Mass spectra of lipids extracted from 1205Lu cells treated with DMOS as control and avasimibe (10 µM, 2 days). m/z 614.6, m/z 640.6, m/z 642.6, m/z 668.6, and m/z 670.7 stand for cholesteryl myristate (C14:0), cholesteryl palmitoleate (C16:1), cholesteryl palmitate (C16:0), cholesteryl oleate

(C18:1), and cholesteryl stearate (C18:0), respectively. The spectral intensity shown in (a) and (c) was normalized by the $CH_2$ bending band at 1445 $cm^{-1}$.

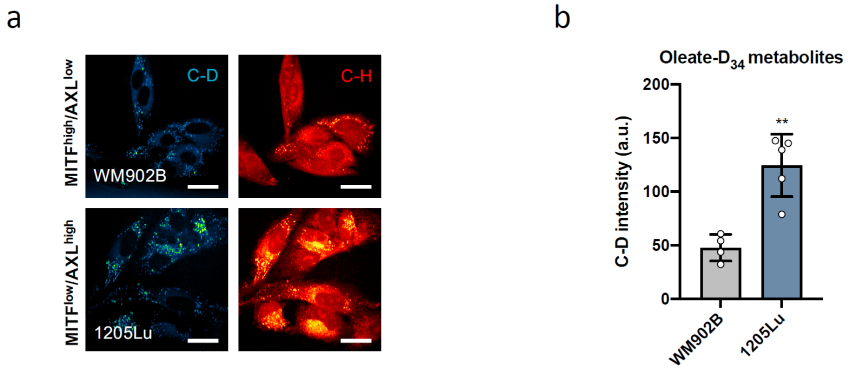


**Supplementary Figure 8. MITF$^{low}$/AXL$^{high}$ melanoma shows higher oleate uptake activity compared to MITF$^{high}$/AXL$^{low}$ melanoma.** (a) Representative SRS images in the C-D (2127 cm$^{-1}$) and C-H (2899 cm$^{-1}$) regions of WM902B and 1205Lu, cultured with oleate-$D_{34}$ containing media (100 µM, 6 hours). (b) Quantification of SRS intensity at 2127 cm$^{-1}$ from C-D positive LDs. Scale bars, 10 µm. Data represent mean ± SD. **: $p < 0.01$.

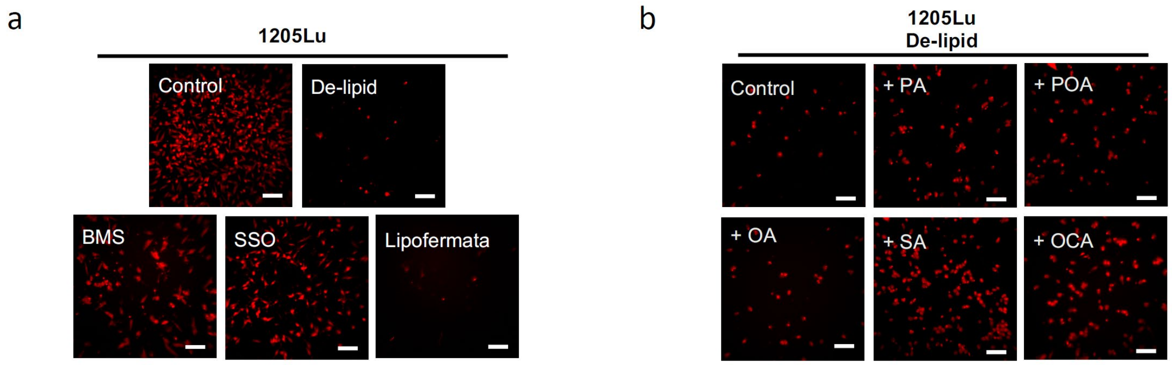


**Supplementary Figure 9. Fatty acid sapienate significantly promotes cell migration.** (a) Images of migrated 1205Lu pre-cultured with de-lipidized medium or pre-treated with BMS309403 (BMS, 50 µM, 1 day), sulfosuccinimidyl oleate (SSO, 50 µM, 1 day) and lipofermata (10 µM, 1 day). (b) Images of migrated 1205Lu pre-cultured with de-lipidized serum media (1 day) and supplemented with ethanol as control and fatty acids (20 µM, 12 hours) as indicated. Control group was used for normalization. Scale bars, 50 µm.

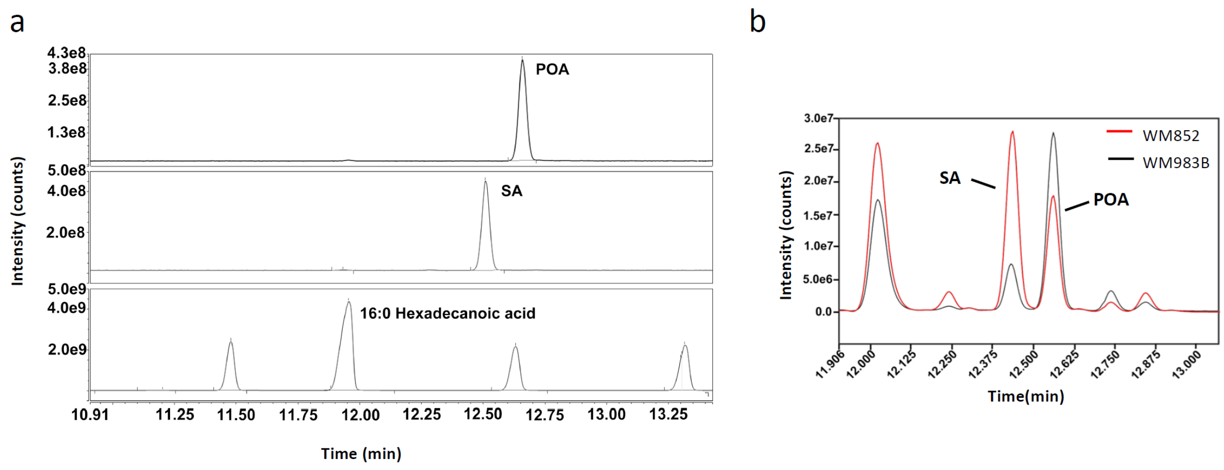


**Supplementary Figure 10.** (a) Detection of sapienate (SA) and palmitoleate (POA) by GC/MS. 16:0 Hexadecanoic acid was used as an internal control. (b) Mass spectra of lipids extracted from WM852 and WM983B. The peaks at 12.38 min and at 12.61 min stand for sapienate (SA) and palmitoleate (POA), respectively.

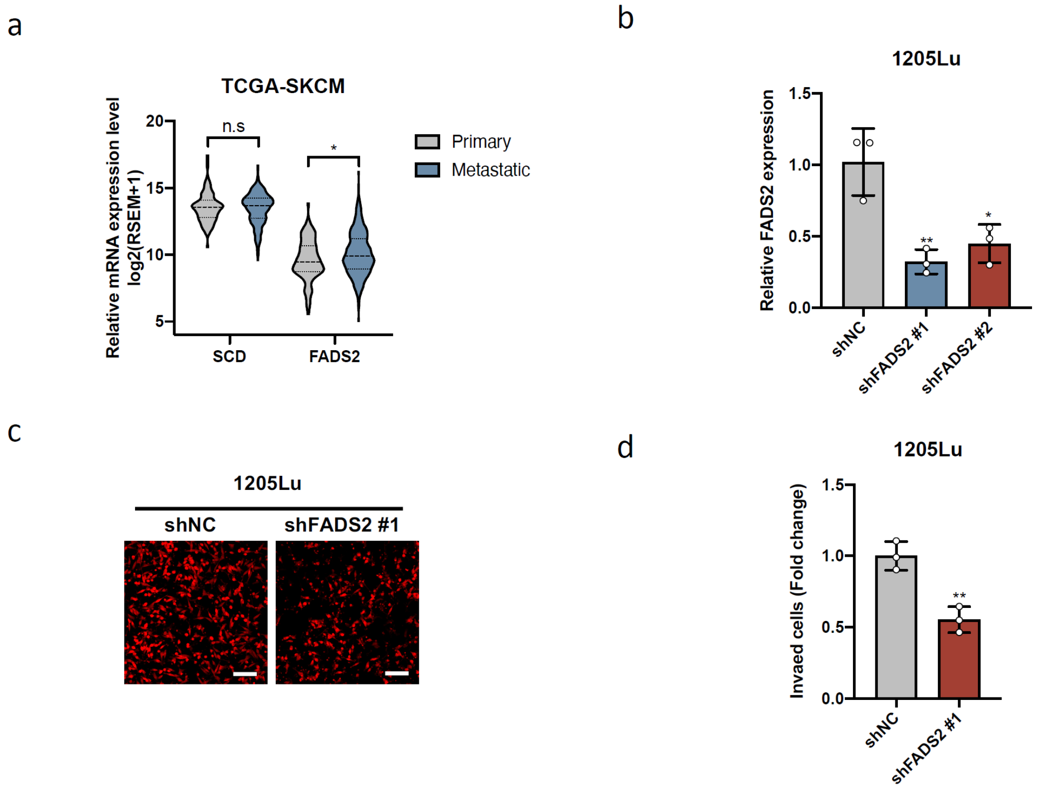


**Supplementary Figure 11. Inhibition of FADS2 suppresses melanoma invasion.** (a) Relative mRNA expression levels of SCD and FADS2 in primary and metastatic melanoma from TCGA-SKCM database. (b) Relative FADS2 mRNA levels in 1205Lu stabling expressing control shRNA (shNC) or FADS2 shRNAs (shFADS2 #1 and #2). (c) Images and (d) quantification of invaded 1205Lu cells stabling expression shNC and shFADS2. Scale bars, 50 µm. Data represent mean ± SEM. *: $p < 0.05$, **: $p < 0.01$, n.s.: not significant.

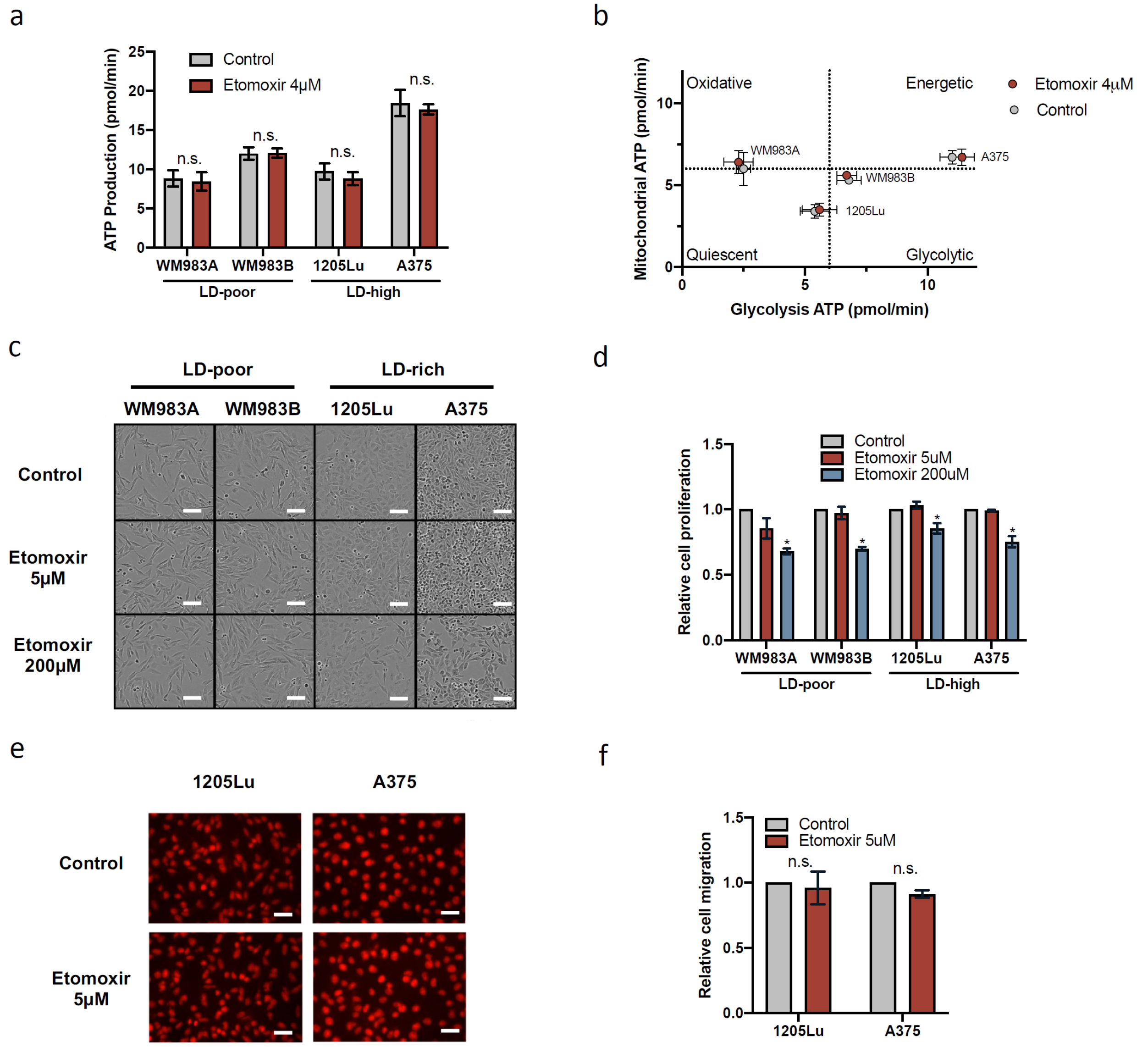


**Supplementary Figure 12. Fatty acid β-oxidation is not the major source of energy in melanoma.** (a) ATP production in LD-poor and LD-rich melanoma cells treated with ethanol as control and 4 μM etomoxir. (b) A plot of mitochondrial ATP versus glycolysis ATP productions in melanoma cells treated with ethanol as control and 4 μM etomoxir. (c) Brightfield images of melanoma cells treated with ethanol as control, 5 μM etomoxir, and 200 μM etomoxir. Scale bar: 50 μm. (d) Quantification of cell proliferation in melanoma cells. (e) Images and (f) quantification of migrated 1205Lu and A375 pre-treated with ethanol as control and 5 μM etomoxir. Scale bar: 25 μm. Data present mean ± SD. *: $p < 0.05$, n.s.: not significant.

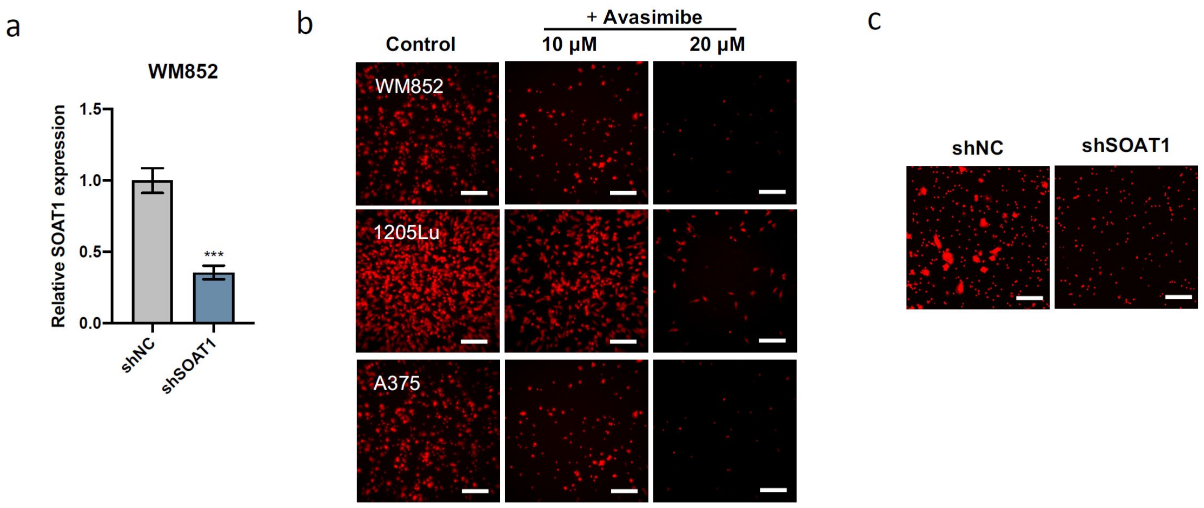


**Supplementary Figure 13. Inhibition of cholesterol esterification suppresses melanoma migration.** (a) Relative SOAT1 mRNA levels in WM852 stabling expressing control shRNA (shNC) or SOAT1 shRNAs (shSOAT1). (b) Images of migrated melanoma cells treated with DMSO as control, 10 µM avasimibe and 20 µM avasimibe for 2 days. (c) Images of migrated WM852 expressing shNC and shSOAT1. Scale bars, 50 µm. Data represent mean ± SEM. ***: $p < 0.001$.

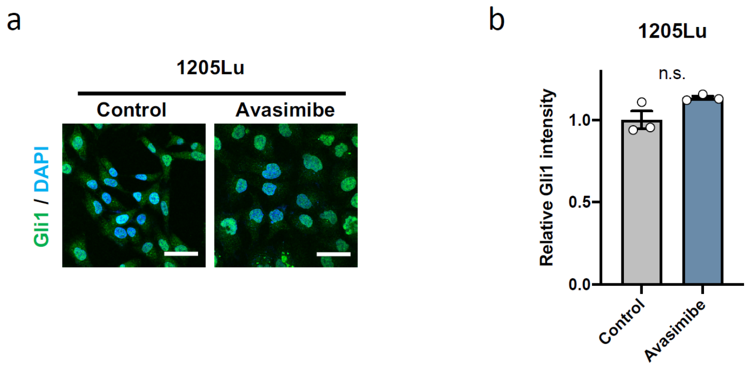


**Supplementary Figure 14. Inhibiting cholesterol esterification does not affect Gli1 nucleus localization.** (a) Fluorescence images of immunostaining Gli1 in 1205Lu treated with DMSO as control and avasimibe (10 µM, 2 days). Scale bars, 25 µm. (b) Quantification of nuclear Gli1 in 1205Lu as shown in (a). Data represent mean ± SD. n.s.: not significant.

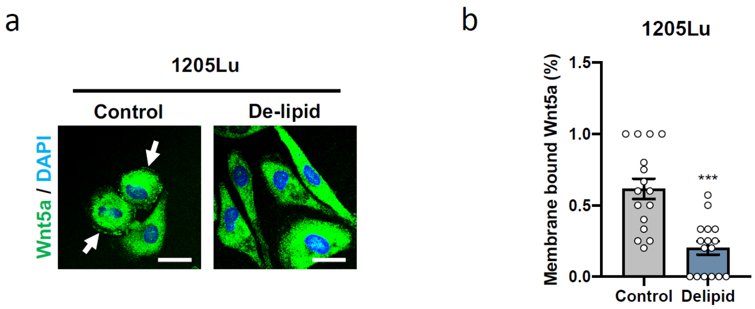


**Supplementary Figure 15. Suppressing fatty acid uptake reduces membrane bound Wnt5a.** (a) Fluorescence images of immunostaining Wnt5a in 1205Lu cultured with de-lipidized serum for 2 days. Arrows indicate membrane bound Wnt5a. Scale bars, 25 μm. (b) Quantification of membrane bound Wnt5a in 1205Lu as shown in (a). Data represent mean ± SD. ***: $p < 0.001$.
